# *cis*-Regulatory Chromatin Contacts in Neural Cells Reveal Contributions of Genetic Variants to Complex Neurological Disorders

**DOI:** 10.1101/494450

**Authors:** Michael Song, Xiaoyu Yang, Xingjie Ren, Lenka Maliskova, Bingkun Li, Ian Jones, Chao Wang, Fadi Jacob, Kenneth Wu, Michela Traglia, Tsz Wai Tam, Kirsty Jamieson, Si-Yao Lu, Guo-Li Ming, Jun Yao, Lauren A. Weiss, Jesse Dixon, Luke M. Judge, Bruce R Conklin, Hongjun Song, Li Gan, Yin Shen

## Abstract

Mutations in gene regulatory elements have been associated with a wide range of complex neurological disorders. However, due to their inherent cell type-specificity and difficulties in characterizing their regulatory targets, our ability to identify causal genetic variants has remained limited. To address these constraints, we perform integrative analysis of chromatin interactions using promoter capture Hi-C (pcHi-C), open chromatin regions using ATAC-seq, and transcriptomes using RNA-seq in four functionally distinct neural cell types: iPSC-induced excitatory neurons and lower motor neurons, iPSC-derived hippocampal dentate gyrus (DG)-like neurons, and primary astrocytes. We identify hundreds of thousands of long-range *cis* interactions between promoters and distal promoter-interacting regions (PIRs), enabling us to link regulatory elements to their target genes and reveal putative pathways that are dysregulated in disease. We validate several novel PIRs using CRISPR techniques in human excitatory neurons, demonstrating that *CDK5RAP3, STRAP*, and *DRD2* are transcriptionally regulated by physically linked enhancers. Finally, we show that physical chromatin interactions mediate genetic interactions in autism spectrum disorder (ASD). Our study illustrates how characterizing the 3D epigenome elucidates novel regulatory relationships in the central nervous system (CNS), shedding light on previously unknown functions for noncoding variants in complex neurological disorders.

## Introduction

A large number of genetic variations associated with diverse human traits and diseases are located in putative regulatory regions. Genetic lesions in these regulatory elements can contribute to complex human disease by modulating gene expression and disrupting finely tuned transcriptional networks in development and function. However, deciphering the roles of regulatory variants in disease pathogenesis remains nontrivial due to their lack of annotation in the physiologically relevant cell types. Furthermore, regulatory elements often interact with their cognate genes over long genomic distances, precluding a straightforward mapping of regulatory element connectivity and limiting the functional interpretation of noncoding variants from genome wide association studies (GWAS). Typically, neighboring genes are assigned as risk loci for noncoding variants. However, this nearest gene model is challenged both by experimental and computational evidence^1,2^. For instance, two independent obesity-associated SNPs in the *FTO* gene have been shown not to regulate *FTO*, but *IRX3* in the brain and both *IRX3* and *IRX5* in adipocytes, respectively^3,4^. The *FTO* locus in obesity illustrates the potentially intricate and cell type-specific manner in which noncoding variants contribute to disease. However, such well-annotated cases are rare, and we still lack systematic mappings of GWAS SNPs to their regulatory targets, especially in the context of complex neurological disorders.

Previous epigenomic annotations of the germinal zone (GZ) and cortical and subcortical plates (CP) in the human brain revealed the importance of 3D chromatin structure in gene regulation and disease^5,6^. However, these studies utilized complex, heterogeneous tissues, limiting their abilities to interpret gene regulation in a cell type-specific manner. Therefore, charting the landscape of epigenomic regulation in well-characterized, physiologically relevant cell types should offer significant advantages for identifying causal variants, deciphering their functions, and enabling novel therapies for previously intractable diseases. Towards this goal, we used wild type human induced pluripotent stem cells (iPSCs) from the WTC11 line^7^ to generate three neuronal cell types: excitatory neurons^8^, hippocampal dentate gyrus (DG)-like neurons^9^, and lower motor neurons^10^. GFAP-positive astrocytes from the gastrulating brains of two individuals were also included for their relevance to human brain development and disease. By performing integrative analysis of promoter-centric, long-range chromatin interactions, open chromatin regions, and transcriptomes (**Fig. 1a**), we provide comprehensive annotations of promoters and distal promoter-interacting regions (PIRs) for each of the neural cell types. We identify putative gene targets for both *in vivo* validated enhancer elements from the VISTA Enhancer Browser^11^ and disease-associated variants, enabling the functional validation of PIRs driving diverse processes in cellular identity and disease.

**Figure 1.**
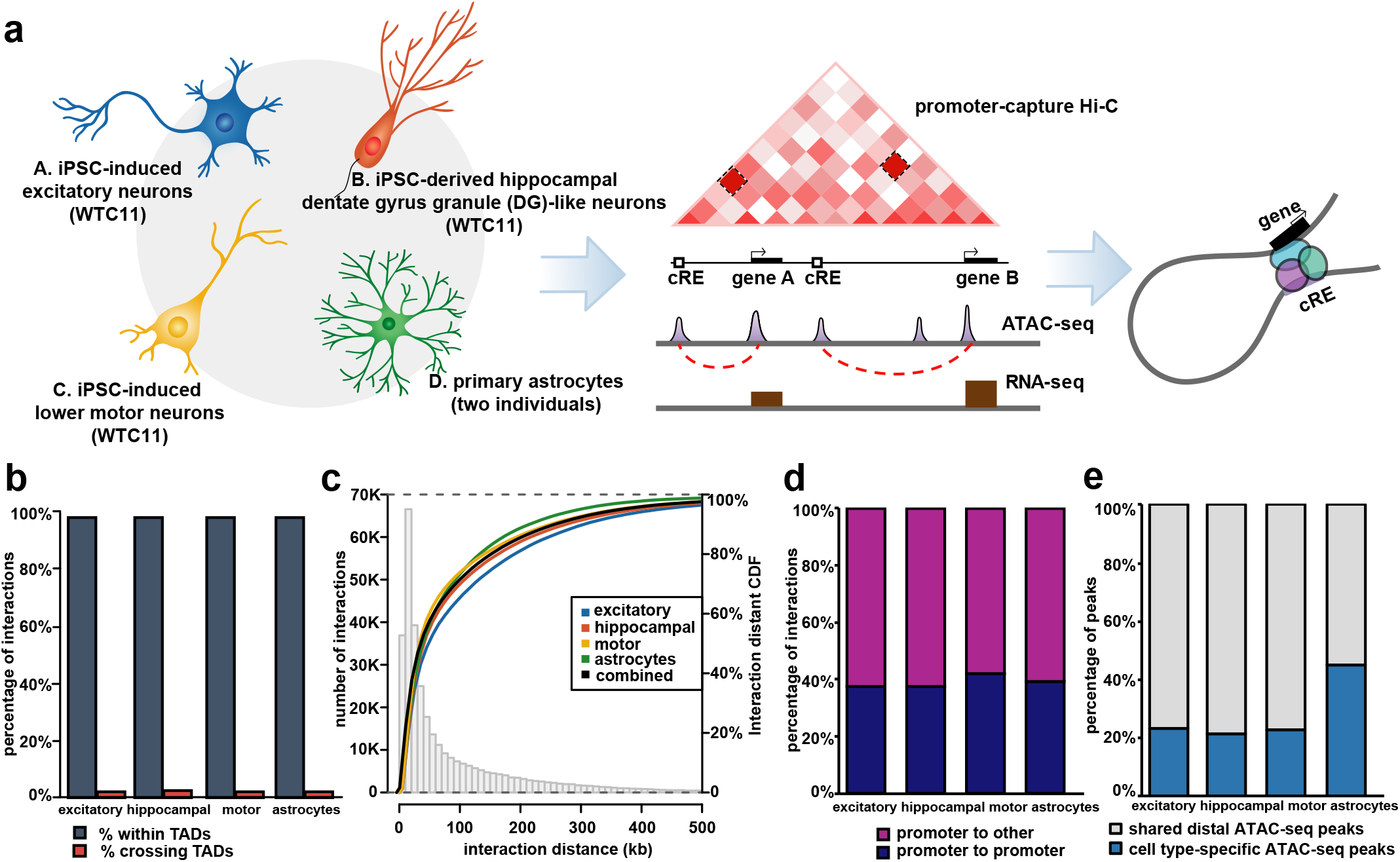
Genome-wide mapping of physical chromatin interactions in functionally distinct neural cell types. (**a**) Schematic of the study design for generating four functionally distinct cell types in the CNS and performing integrative analysis of chromatin interactions using promoter capture Hi-C, open chromatin regions using ATAC-seq, and transcriptomes using RNA-seq. For pcHi-C, we used 3, 2, 3, and 4 biological replicates respectively for the excitatory neurons, hippocampal DG-like neurons, lower motor neurons, and astrocytes. For ATAC-seq, we used 2, 2, 3, and 4 biological replicates respectively for the cell types. For RNA-seq, we used 2, 2, 2, and 4 biological replicates respectively for the cell types. (**b**) Proportions of interactions occurring within TADs for each cell type. (**c**) Histogram and empirical CDF plots of interaction distances for each cell type. (**d**) Proportions of interactions between promoter-containing bins (blue) and promoter- and non-promoter-containing bins (purple) for each cell type. (**f**) Proportions of cell type-specific (blue) and shared (grey) distal open chromatin peaks at PIRs for each cell type.

## Results

### Characterizing the epigenomic landscape of long-range chromatin interactions in human neural cells

To investigate general features of the epigenomic landscape for specific cell types in the human central nervous system (CNS), we focused on isogenic iPSC-induced excitatory neurons, iPSC-derived hippocampal dentate gyrus (DG)-like neurons, and iPSC-induced lower motor neurons, three neuronal subtypes which are currently impractical to isolate from primary tissue. Excitatory neurons were induced from a wild type male iPSC line (WTC11) containing an integrated, isogenic, and inducible neurogenin-2 (Ngn2) cassette (i^3^N iPSCs) with doxycycline-inducible *Ngn2* at the AAVS1 safe-harbor locus^8^. The i^3^N iPSCs enabled us to obtain homogenous excitatory neurons expressing both the cortical neuron marker CUX1 and the glutamatergic neuron marker VGLUT1^8,12^ (**Supplementary Fig. 1a, b**).

Hippocampal DG-like neurons expressing the DG granule cell marker PROX1 were differentiated from a WTC11 iPSC line using defined factors as described in previous publications^9,13^ (**Supplementary Fig. 1a, b**). Finally, lower motor neurons were induced from a WTC11 line containing integrated, isogenic, and inducible copies of *NGN2, ISL1*, and *LHX3* at the AAVS1 safe-harbor locus (i^3^LMN iPSCs)^10^. The cells exhibited homogenous expression of the lower motor neuron markers HB9 and SMI32 in culture (**Supplementary Fig. 1a, b**). In addition, all three neuronal subtypes exhibited high expression of the synaptic genes *SYN1* and *SYN2*, the NMDA receptor genes *GRIN1* and *GRIN2A*, and the AMPA receptor genes *GRIA1* and *GRIA2*, evidencing mature synaptic functions (**Supplementary Fig. 1b**). We included two batches of astrocytes isolated from 19 week gastrulating male fetal brain samples using GFAP as a selection marker (ScienCells). Astrocytes were cultured for two or fewer passages *in vitro* and confirmed for positive expression of GFAP prior to harvesting (**Supplementary Fig. 1a**). Based on the age of the donors and transcriptional signatures for dozens of marker genes distinguishing astrocyte progenitor cells (APCs) and mature astrocytes (e.g. high expression of the APC markers *TOP2A* and *TNC* and low expression of the mature astrocyte markers *AGXT2L1* and *WIF1*)^14^, the astrocytes were determined to most likely be APCs (**Supplementary Fig. 1b**).

We constructed pcHi-C, ATAC-seq, and RNA-seq libraries using two to four biological replicates for each cell type (**Supplementary Table 1**). Specifically, promoter-centric, long-range chromatin interactions were mapped using a set of 280,445 unique RNA probes targeting the promoters of 19,603 coding genes in GENCODE 19 (Jung et al., in revision). We first confirm the reproducibility of contact frequency and saturation of inter-replicate correlation for our pcHi-C libraries using HiCRep (**Supplementary Fig. 2c, d**). Hierarchical clustering of ATAC-seq read density and gene expression similarly group the replicates by cell type (**Supplementary Fig. 2a, b**), evidencing minimal variations during the cell derivation process. Using CHiCAGO^15^, we identified significant chromatin interactions with score ≥ 5 at a total of 195,322 unique interacting loci across the four cell types, with 73,890, 108,156, 66,978, and 84,087 significant interactions being represented in the excitatory neurons, hippocampal DG-like neurons, lower motor neurons, and astrocytes, respectively (**Supplementary Table 2**). Overall, 17,065 or 83.9% of coding gene promoters participate in interactions in at least one cell type (**Supplementary Fig. 1c**), with 80% of PIRs interacting within a distance of 160 kb (**Fig 1c, Supplementary Fig. 3a**). Furthermore, over 97% of interactions reside within topologically associating domains (TADs) from Hi-C datasets in human fetal brain tissue^6^ (**Fig. 1b**). Approximately 40% of interactions occur between promoter-containing bins, while the remaining 60% occur between promoter- and non-promoter-containing bins (**Fig. 1d**). The observed numbers of promoter-promoter interactions can potentially be attributed to transcriptional factories of co-regulated genes, widespread co-localization of promoters^16,17^, and the ability of many promoters to doubly function as enhancers^18,19^. Finally, up to 40% of interacting distal open chromatin peaks are specific to each cell type (**Fig. 1e**), suggesting that PIRs are capable of orchestrating cell type-specific gene regulation. Astrocytes in particular exhibit the largest proportion of cell type-specific open chromatin peaks, likely reflecting basic differences between the neuronal and glial lineages.

We observe that the majority of promoters interact with more than one PIR (**Fig. 2a**). This observation is consistent with the large number of regulatory elements in the human genome^20^ and previous findings that each promoter can be regulated by multiple enhancers^21^. To examine global chromatin signatures at PIRs, we leveraged chromatin states inferred by ChromHMM^22^ in matched human brain tissues from the Roadmap Epigenomics Project^23^ (dorsolateral prefrontal cortex for excitatory neurons, hippocampus middle for hippocampal DG-like neurons, and normal human astrocytes for fetal astrocytes). We show that PIRs are highly enriched for active chromatin features including open chromatin peaks, enhancers, and transcriptional start sites (TSSs) while simultaneously exhibiting depletion for repressive heterochromatin marks (**Fig. 2b**). PIRs are also enriched for H3K27ac and CTCF binding sites mapped using CUT&RUN^24^ in excitatory neurons and lower motor neurons, as well as ChIP-seq in astrocytes from ENCODE (**Supplementary Fig. 3b**). We find that promoters interacting exclusively with enhancer-PIRs exhibit elevated levels of transcription compared to promoters interacting exclusively with repressive-PIRs (two sample t-test, one-sided, *p*=9.4×10^−63^) (**Fig. 2c, Supplementary Fig. 3c**). Since multiple enhancers can interact with and regulate the same promoter, we assessed whether interactions with multiple enhancer-PIRs could present evidence for additive effects on transcription. By grouping genes according to the number of interactions their promoters form with enhancer-PIRs in each cell type, we discover a positive correlation between the number of enhancer-PIR interactions and the mean gene expression in each group (linear regression test, *p*=2.1×10^−3^) (**Fig. 2d, Supplementary Fig. 3d**). Our results demonstrate that chromatin interactions enable the identification of PIRs which are not only enriched for regulatory features, but which can also modulate gene expression.

**Figure 2.**
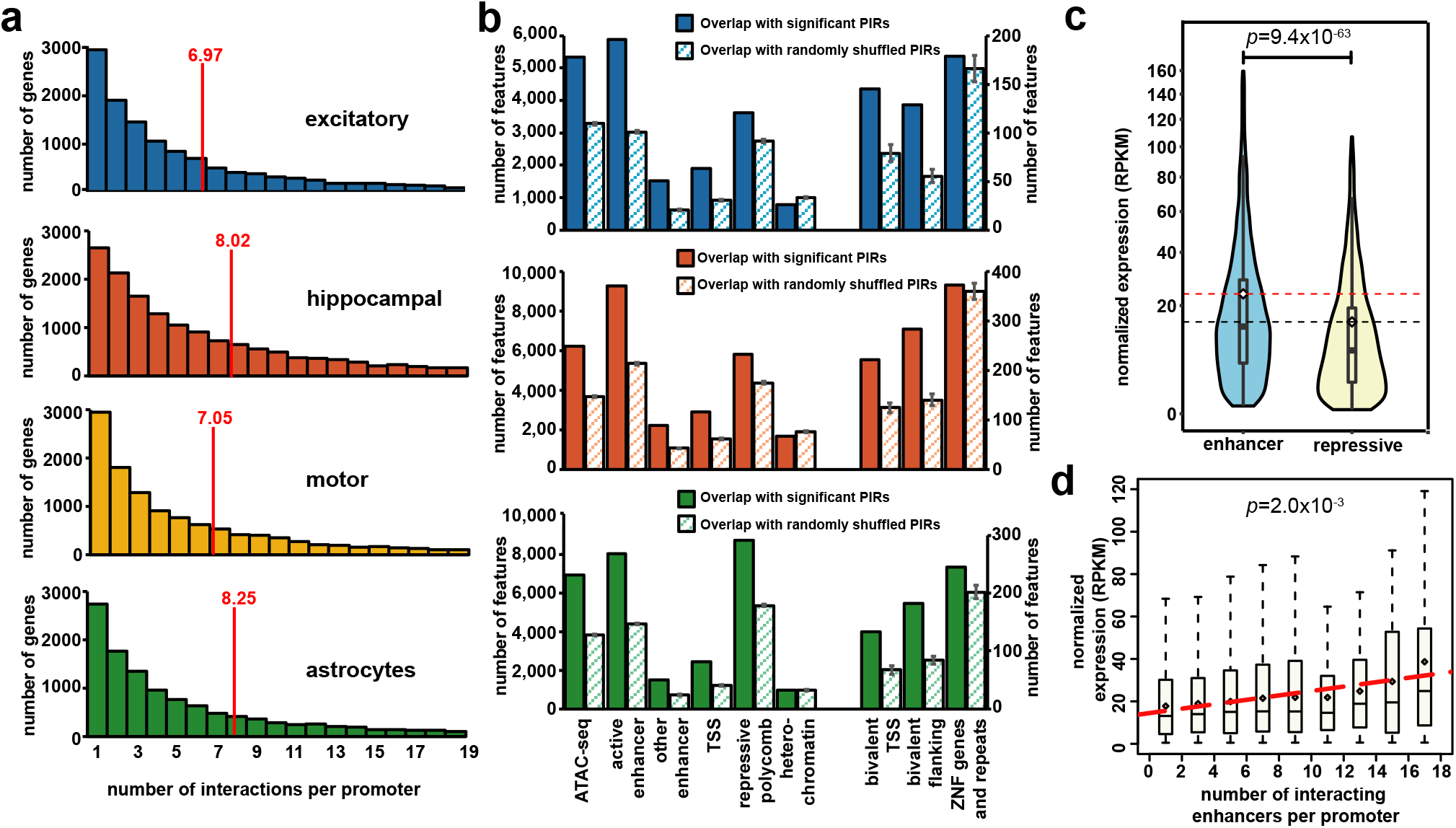
Integrative analysis of chromatin interactions, epigenomic features, and gene expression. (**a**) Histograms of the numbers of PIRs interacting with each promoter in each cell type. The means are indicated with red lines. Only promoters interacting with at least one PIR are included (15,316 promoters in excitatory neurons, 19,546 promoters in hippocampal DG-like neurons, 14,990 promoters in lower motor neurons, and 15,397 promoters in astrocytes, out of a total of 34,401 protein coding and noncoding RNA promoters in GENCODE 19). (**b**) Bar plots showing counts of epigenomic chromatin states (inferred at a 200 bp resolution using the ChromHMM core 15 state model in matched tissues) overlapping significant (solid bars) versus randomly shuffled (striped bars) PIRs for each cell type. Error bars represent the standard deviation over 100 sampled sets of randomly shuffled PIRs. No matching tissue data was available for the lower motor neurons so they were omitted from the analysis. (**c**) Comparative gene expression analysis across all cell types for expressed genes (normalized RPKM > 0.5) whose promoters interact exclusively with either enhancer-PIRs (n=6836) or repressive-PIRs (n=2612). Distributions of gene expression values are shown for each group. (**d**) Boxplots showing distributions of gene expression values across all cell types for expressed genes (normalized RPKM > 0.5) grouped according to the numbers of interactions their promoters form with enhancer-PIRs. Linear regression was performed on the mean gene expression values for each bin. Only bins containing at least 10 genes were included in the analysis.

### PIRs contribute to cellular identity

We find that chromatin interactions exhibit distinct patterns of cell type specificity, with tens of thousands of interactions that are specific to each cell type (**Fig. 3a, Supplementary Fig. 4a**). These interactions may underlie important functional differences between the cell types, as gene ontology (GO) enrichment analysis for genes interacting with cell type-specific PIRs produced terms associated with neuronal function in the neuronal subtypes and immune function in the astrocytes (**Fig. 3b, Supplementary Table 3**). Meanwhile, 58,809 or 30.1% of unique interactions are shared across all four cell types, with neural precursor cell proliferation and neuroblast proliferation ranking among the top GO terms for genes participating in shared interactions. In conjunction with the observed enrichment of active chromatin signatures at PIRs, the cell type-specific nature of PIRs suggests that they harbor lineage-specific regulatory roles. Indeed, numerous promoters of differentially expressed genes form specific contacts with PIRs in the corresponding cell types, including *OPHN1* in hippocampal DG-like neurons, *CHAT* in lower motor neurons, and *TLR4* in astrocytes (**Supplementary Fig. 4b**). OPHN1 is known to stabilize synaptic AMPA receptors and mediate long-term depression in the hippocampus, and its loss of function is associated with X-linked mental retardation^25^. Meanwhile, CHAT is a principal marker for lower motor neuron maturity and function, and TLR4 is a key regulator of immune activation and synaptogenesis in astrocytes^26^. Finally, hierarchical clustering of interaction scores across all significant promoter-PIR interactions demonstrates that cell types can reliably be grouped according to lineage-specific features, with the three neuronal subtypes clustering together more tightly than with the astrocytes (**Fig. 3a**).

**Figure 3.**
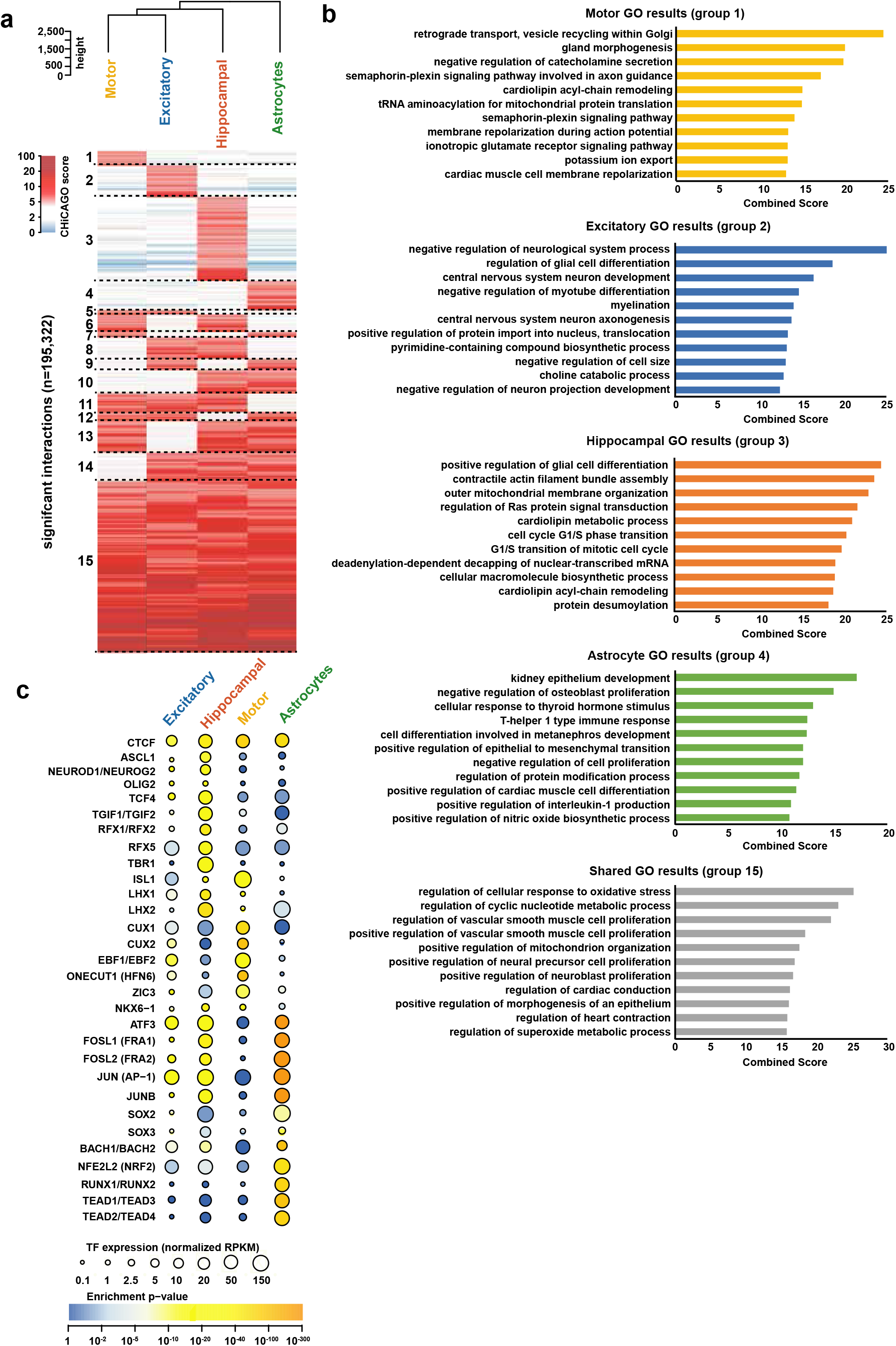
Cell type-specific PIRs and TF motif enrichment analysis. (**a**) Classification of unique promoter-PIR interactions with interaction score ≥ 5 in at least one cell type into specificity categories based on their scores in each cell type. The numbers of promoter-PIR interactions in each specificity category are summarized in **Supplementary Fig. 3a**. Cell types are also hierarchically clustered based on their interaction scores over all interacting loci. (**b**) Top enriched GO terms from the “GO Biological Process 2018” ontology in Enrichr for genes whose promoters participate in cell type-specific interactions with distal open chromatin peaks in each cell type (groups 1-4). Also shown are top enriched GO terms for genes participating in shared interactions across all cell types (group 15). Enriched GO terms are ranked by their combined scores (calculated by multiplying the log of the p-value by the z-score of the deviation from the expected rank). An expanded list of enriched GO terms and their raw p-values is available in **Supplementary Table 3**. (**c**) Enrichment of consensus TF motif sequences at open chromatin peaks in cell type-specific PIRs using HOMER, organized by motifs (rows) and cell types (columns). The color of each dot represents the degree of enrichment (negative log p-value) for each motif in each cell type, while the size of each dot represents gene expression (normalized RPKM) for the corresponding TFs for each motif. Entries with similar or identical consensus TF motif sequences are grouped for brevity.

Gene expression is coordinately controlled by transcription factors (TFs) and regulatory elements such as enhancers. Therefore, PIRs identified through chromatin interactions provide a unique perspective to investigate potential mechanisms underlying cell type-specific gene regulation. We use HOMER^27^ to evaluate TF motif enrichment at cell type-specific distal open chromatin peaks in PIRs for each cell type (**Fig. 3c, Supplementary Table 4**). We find that the CTCF motif is highly enriched across all cell types, consistent with its role in mediating looping between promoters and regulatory elements within TADs^28–31^. Furthermore, motifs for ASCL1, ISL1, NEUROG2, OLIG2, and ZIC3, TFs linked to neuronal fate commitment, are enriched in various patterns across the neuronal subtypes. Other TFs functioning in brain development include CUX1/CUX2, EBF1/EBF2, HFN6, LHX1/LHX2, NKX6-1, TCF4, TGIF2, and the RFX factors. The TBR1 motif is enriched in hippocampal DG-like neurons, consistent with TBR1’s roles in NMDA receptor assembly and maintaining long-term potentiation in the hippocampus^32^. Meanwhile, astrocytes are enriched for motifs in the Fos and Jun families, which contain key regulators for inflammatory and immune pathways. Also enriched in astrocytes are motifs for ATF3 and the RUNX and TEAD families, TFs with established roles in astrocyte differentiation, maturation, and proliferation. Motif enrichment is not always accompanied by expression of the corresponding TFs, potentially reflecting synergistic interactions between different cell types in the CNS. For example, NRF2 is a key regulator of the oxidative stress response, and its activity has been shown to be repressed in neurons while inducing a strong response in astrocytes^33^. Therefore, its shared expression may reflect the neuroprotective role that astrocytes provide for other cell types. Alternatively, TFs do not have to be expressed at high levels to perform their cellular functions due to additional avenues for regulation at the post-transcriptional and protein levels. Overall, our results demonstrate that PIRs contribute to cell fate commitment and are capable both of recapitulating known and revealing novel regulators.

### *In vivo* validation of interactions linking enhancer elements to their target genes

Regulation of target genes by enhancers is thought to be mediated by physical chromatin looping. Congruent with this concept, interactions detected by pcHi-C can be used to link enhancers with their target genes. The VISTA Enhancer Browser^11^ is a database containing experimentally validated human and mouse noncoding sequences with enhancer activity. To date, it contains 2,956 tested elements, 1,568 of which exhibit enhancer activity during embryonic development^11^. However, the regulatory targets for these enhancer elements remain largely uncharacterized. To address this knowledge gap, we provide detailed cell type-specific annotations of putative target genes for each enhancer element using our significant promoter-PIR interactions and open chromatin peaks (**Supplementary Table 5**). In total, our interactions recover 589 or 37.6% of positively annotated enhancer elements with human sequences, 320 of which were further annotated as neural enhancers according to tissue-specific patterns of LacZ staining in mouse embryos (**Fig. 4a, b**). Of the 589 interacting positive enhancer elements, only 60 interact exclusively with their nearest genes (scenario III), whereas 306 interact exclusively with more distal genes (scenario I), identifying 464 novel gene targets (**Fig. 4c**). Meanwhile, 118 elements interact with both their nearest genes and a total of 484 more distal genes (scenario II). The remaining 105 elements cannot be resolved at the HindIII fragment level for interactions with their nearest genes (scenario IV), though they interact with 395 additional non-neighboring genes. In total, our chromatin interactions identify 1,343 novel, putative gene targets for positive enhancer elements in the VISTA Enhancer Browser, significantly expanding our knowledge of gene regulatory relationships at these loci.

**Figure 4.**
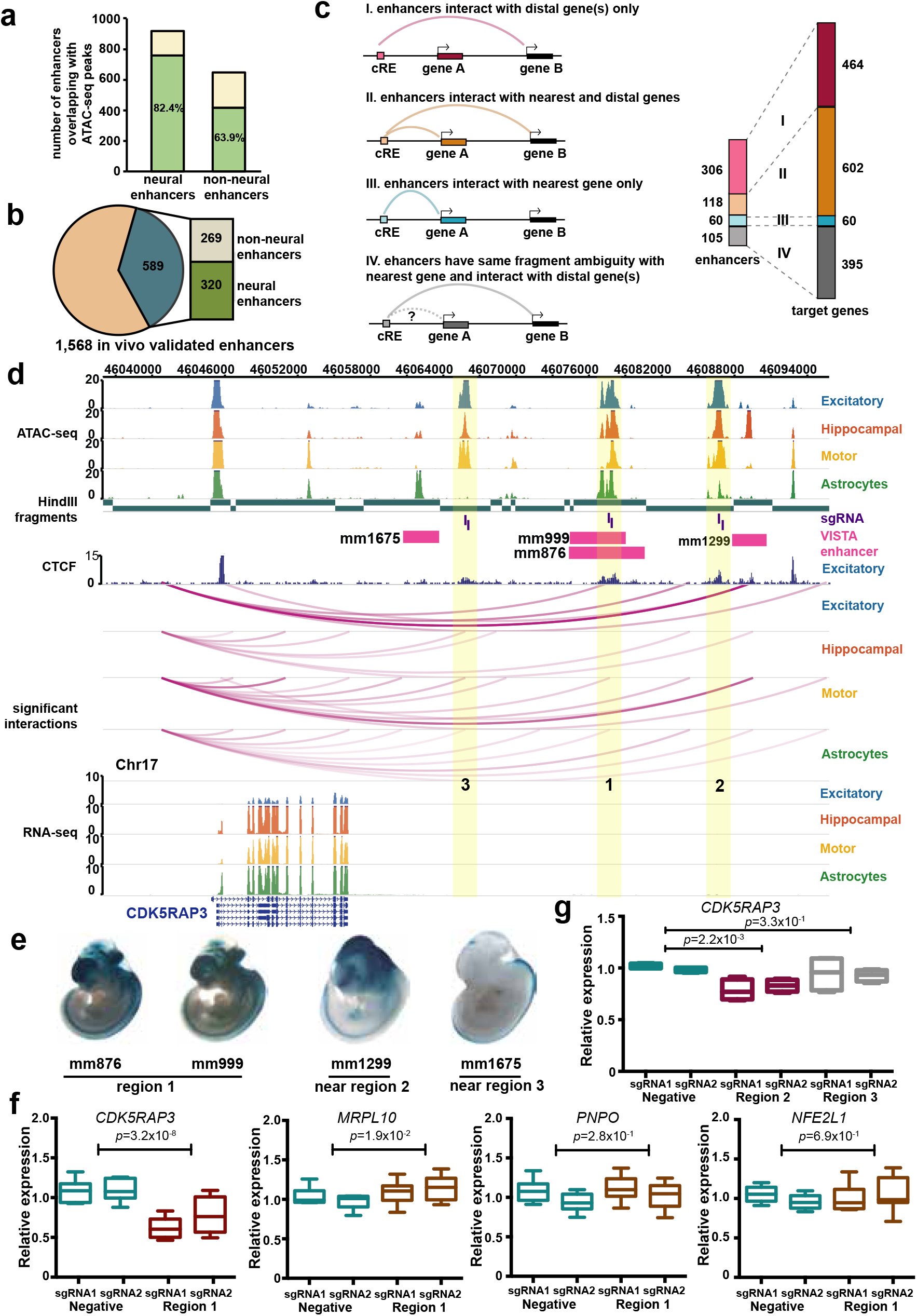
Validation of PIRs in human neural cells. (**a**) *In vivo* validated enhancers with neural annotations overlap a significantly higher proportion of open chromatin peaks in the neural cells compared to enhancers with non-neural annotations (chi-squared test, p<2.2×10^−16^). (**b**) Pie chart showing counts of *in vivo* validated enhancers with human sequences participating in chromatin interactions (589 out of 1568 total elements). Counts of interacting positive enhancer elements with neural and non-neural annotations are also shown. (**c**) Counts of interacting positive enhancer elements interacting exclusively with their nearest genes (blue), interacting exclusively with more distal genes (pink), or interacting with both their nearest genes and more distal genes (orange). Positive enhancer elements that could not be resolved for interactions with their nearest genes are also shown (grey). The number of regulatory targets interacting with positive enhancer elements in each category is shown on the right. (**d**) Promoter-PIR interactions at the *CDK5RAP3* locus. Open chromatin peaks in PIRs up to 40 kb downstream of *CDKRAP3* (regions 1, 2, and 3, yellow highlight) interact with the promoter of *CDK5RAP3* in a cell type-specific manner. Notably, only regions 1 and 2 participate in interactions with the promoter of *CDK5RAP3* in excitatory neurons. In addition, both *in vivo* validated enhancers (pink) and CTCF binding sites in excitatory neurons (dark blue) are shown to be localized to all three candidate regulatory regions. All interactions fall within a CP TAD (chr17:45,920,000-47,480,000). (**e**) LacZ staining in mouse embryos shows tissue-specific patterns of enhancer activity. (**f**) CRISPRi silencing of region 1 using two independent sgRNAs results in significant downregulation of *CDK5RAP3* expression in excitatory neurons (two sample t-test, two-sided, *p*=3.2×10^−8^). No significant downregulation was detected for the neighboring genes *MRPL10, PNPO*, and *NFE2L1*. Each CRISPRi experiment was performed in triplicate, with three technical replicates per experiment. (**g**) CRISPRi silencing of region 2, but not region 3, results in significant downregulation of *CDK5RAP3* expression in excitatory neurons (two sample t-test, two-sided, *p*=2.2×10^−3^).

### Validation of PIRs detected in human neural cells using CRISPR techniques

We performed validation of two PIRs (regions 1 and 2) physically interacting up to 40 kb away with the promoter of *CDK5RAP3* (**Fig. 4d**). CDK5RAP3 is a known regulator of CDK5, which functions in neuronal development^34^ and regulates proliferation in non-neuronal cells^35^. Both PIRs overlap open chromatin peaks as well as enhancer elements with positively annotated forebrain activity in the VISTA Enhancer Browser (mm8766 and mm999 for region 1 and mm1299 for region 2) (**Fig. 4e**). We targeted both regions for CRISPR deletion in i^3^N iPSCs, followed by differentiation of the iPSCs into excitatory neurons and quantification of any changes in gene expression by qPCR. Deleting the 2 kb open chromatin peak in region 1 led to significant downregulation of *CDK5RAP3* expression (two sample t-test, two-sided, *p*=7.7×10^−7^) (**Supplementary Fig. 4c**). Upon trying to delete the open chromatin peak in region 2, we observed massive cell death of iPSCs immediately following introduction of the Cas9-sgRNA protein complex. We picked 48 individual clones from cells surviving the transfection, but failed to isolate any clones with deletions, suggesting that this locus is essential for maintaining *CDK5RAP3* expression and survival in iPSCs. To circumvent the lethal phenotype for iPSCs associated with region 2, we silenced both regions using CRISPR interference (CRISPRi) in excitatory neurons. We also silenced a third region interacting in the other cell types, but not in excitatory neurons (region 3). We show that silencing of regions 1 and 2 but not region 3 leads to significant downregulation of *CDK5RAP3* expression without influencing the expression of nearby genes (two sample t-test, two-sided, *p*=3.2×10^−8^ for region 1 and *p*=2.2×10^−3^ for region 2) (**Fig. 4f-g**). Interestingly, a neighboring enhancer element annotated with spinal cord activity (mm1576) also participates in interactions with the *CDK5RAP3* promoter in lower motor neurons and astrocytes, but not in the excitatory neurons and hippocampal DG-like neurons (**Fig. 4d, e**). These results demonstrate that chromatin interactions recapitulate cell type-specific patterns of enhancer activity, underscoring the importance of studying epigenomic regulation in the appropriate cell types.

### Cell type-specific enrichment and regulatory target identification for complex neurological disorder-associated variants at PIRs

Previous large-scale epigenomic studies of human tissues and cell lines highlighted the importance of disease-associated variants at distal regulatory regions^23^ and the need for high-throughput approaches to prioritize variants for further validation. Therefore, we used our chromatin interactions to annotate complex neurological disorder- and trait-associated variants available from the GWAS Catalog^36^. We downloaded a total of 6,396 unique GWAS SNPs at a significance threshold of 10^−6^ for eleven traits including Alzheimer’s disease (AD), attention deficit hyperactivity disorder (ADHD), autism spectrum disorder (ASD), amyotrophic lateral sclerosis (ALS), bipolar disorder (BD), epilepsy (EP), frontotemporal dementia (FTD), mental processing (MP), Parkinson’s disease (PD), schizophrenia (SCZ), and unipolar depression (UD). The GWAS SNPs were imputed at a linkage disequilibrium (LD) threshold of 0.8 using HaploReg^37^ and filtered to obtain a total of 95,954 unique imputed SNPs across all traits (**Supplementary Table 6**). We find that SNPs are enriched at PIRs in a disease- and cell type-specific manner (**Fig. 5a**), with ASD, MP, and SCZ SNPs enriched at PIRs across all cell types. UD SNPs are exclusively enriched in excitatory neurons and hippocampal DG-like neurons, whereas AD, ADHD, and BD SNPs also exhibit enrichment in lower motor neurons. ALS SNPs are enriched in all the neuronal subtypes but not in astrocytes, consistent with the characterization of ALS as a motor neuron disease and reinforcing evidence for its role in hippocampal degeneration^38^. Notably, PD SNPs are enriched in astrocytes but not in other cell types. The enrichment of PD SNPs at astrocyte-specific PIRs also supports the theory that astrocytes play an initiating role in PD^39^, based on evidence that numerous genes implicated in PD possess functions unique to astrocyte biology, as well as the neuroprotective roles that astrocytes serve for dopaminergic neurons in the substantia nigra. Finally, EP and FTD SNPs are not enriched in any of the cell types, indicating their potential functions in alternative cell types, insufficient study power, or mechanisms acting outside of chromatin-mediated gene regulation.

**Figure 5.**
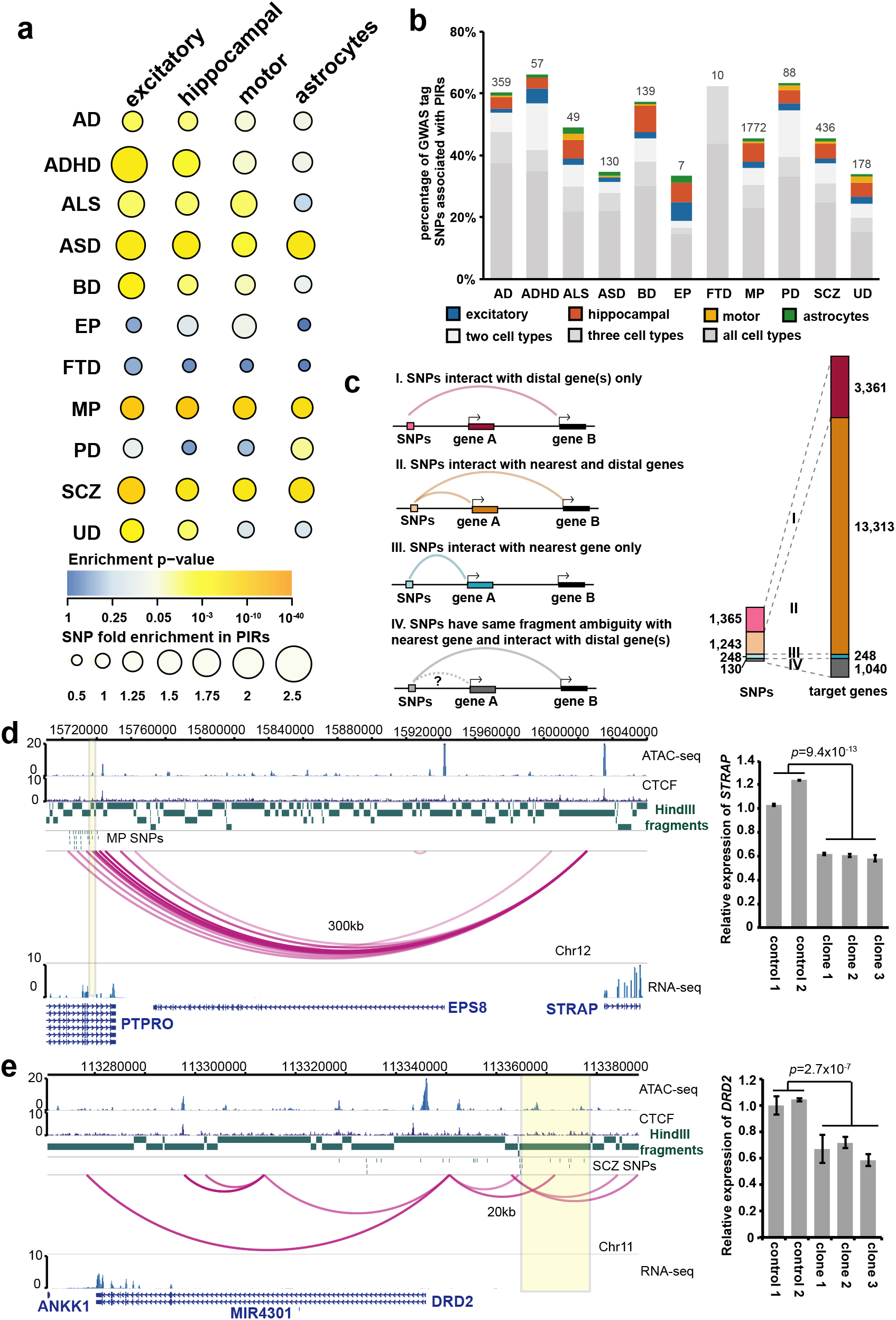
Genetic analysis of promoter-PIR interactions with complex neurological disorder-associated variants. (**a**) Enrichment analysis for complex neurological disorder-associated SNPs in Alzheimer’s disease (AD), attention deficit hyperactivity disorder (ADHD), amyotrophic lateral sclerosis (ALS), autism spectrum disorder (ASD), bipolar disorder (BP), epilepsy (EP), frontotemporal dementia (FTD), mental processing (MP), Parkinson’s disease (PD), schizophrenia (SCZ), and unipolar depression (UD). The color and size of each dot respectively represent the enrichment p-value and raw fold enrichment (calculated as the number of SNPs overlapping significant PIRs divided by the mean number of SNPs overlapping randomly shuffled PIRs across 100 sampled sets) for each cell type and disease pairing. (**b**) Proportions and total counts of GWAS SNPs with at least one SNP in linkage participating in chromatin interactions. Cell type-specific SNPs for excitatory neurons (blue), hippocampal DG-like neurons (orange), lower motor neurons (yellow), and astrocytes (green) are highlighted. (**c**) Counts of GWAS SNPs across all diseases with at least one SNP in linkage interacting exclusively with their nearest genes (scenario III, blue), interacting exclusively with more distal genes (scenario I, pink), or interacting with both their nearest genes and more distal genes (scenario II, orange). GWAS SNPs that could not be resolved for interactions with their nearest genes are also shown (scenario IV, grey). Counts of regulatory targets interacting with GWAS SNPs in each scenario are shown on the right. (**d**) PIRs containing MP SNPS (yellow highlight) in an intron for *PTPRO* interact with the promoter of *STRAP* over 300 kb away. All interactions fall within a CP TAD (chr12:14,960,000-16,040,000). Homozygous deletion of this PIR in three independent clones results in significant downregulation of *STRAP* expression in excitatory neurons (two sample t-test, two-sided, *p*=9.4×10^−13^). Error bars represent the SEM. (**e**) A PIR containing SCZ SNPs interacts with the *DRD2* promoter 20 kb upstream of the PIR. All interactions fall within a CP TAD (chr11:113,200,000-114,160,000). Mono-allelic deletion of this PIR in three independent clones results in significant downregulation of *DRD2* expression in excitatory neurons (two sample t-test, two-sided, *p*=2.7×10^−7^). Error bars represent the SEM.

Up to 70% of GWAS SNPs have at least one SNP in linkage overlapping PIRs in one or more cell types (**Fig. 5b**). As it is common practice in association studies to assign GWAS SNPs to their nearest genes, we evaluated the number of GWAS SNPs with at least one SNP in linkage interacting with their nearest genes. Overall, across all diseases, we find that 248 GWAS SNPs interact exclusively with their nearest genes (scenario III), 1,365 GWAS SNPs interact exclusively with more distal genes (scenario I), and 1,243 GWAS SNPs interact with both their nearest genes and more distal genes (scenario II) (**Fig. 5c, Supplementary Fig. 5a**). Our interactions identify a total of 16,471 non-neighboring gene targets across all diseases (**Supplementary Table 7**). To prioritize variants potentially disrupting regulatory activity, we focused on SNPs overlapping open chromatin peaks at PIRs. We find that PIRs for these putative regulatory SNPs interact with genes possessing functions that are relevant in the context of their respective disease etiologies. For example, GO enrichment results for genes targeted by AD SNPs include terms associated with amyloid beta formation, interferon beta production, and cranial nerve development (**Supplementary Fig. 6, Supplementary Table 9**). Meanwhile, genes targeted by ASD, BD, SCZ, and UD SNPs are enriched for epigenetic terms including chromatin assembly, nucleosome assembly, and nucleosome organization. For genes targeted by SNPs in the remaining diseases, enriched terms include neuronal processes such as myelin maintenance, neuron projection extension, synapse assembly, synaptic transmission, and nervous system development. A comprehensive annotation of PIRs overlapping putative regulatory SNPs is available in **Supplementary Table 8**. Notably, a previously reported interaction between the *FOXG1* promoter and a PIR containing SCZ SNPs over 700 kb away is recapitulated in our chromatin interactions^6^ (**Supplementary Fig. 5b**). In another example, an astrocyte-specific PIR containing AD SNPs targets the promoters of both *CASP2*, a well-known mediator of apoptosis that is also associated with neurodegeneration^40,41^, and *FAM131B*, a putative neurokine (**Supplementary Fig. 7a**). Elsewhere, hippocampal DG-like neuron-specific PIRs containing ASD SNPs target the promoter of *BCAS2*, whose knockdown in mice leads to microcephaly-like phenotypes with reduced learning, memory, and DG volume^42^ (**Supplementary Fig. 7c**). Finally, the *MSI2* promoter is simultaneously targeted by PIRs containing BD SNPs in hippocampal-DG like neurons, lower motor neurons, and astrocytes, as well as a PIR in astrocytes containing SCZ SNPs (**Supplementary Fig. 7d**). Overall, we show that an approach utilizing epigenomic annotations to jointly prioritize variants and identify their regulatory targets enables the identification of regulatory mechanisms with consequential roles in development and disease.

### Validation of PIRs containing neurological disorder-associated regulatory variants

We used CRISPR techniques to validate two PIRs containing putative regulatory SNPs targeting the promoters of the *STRAP* and *DRD2* genes. At the *STRAP* locus, PIRs containing MP SNPs in an intron for *PTPRO* interact over 300 kb away with the promoter of *STRAP* (**Fig. 5d**). STRAP influences the cellular distribution of the survival of motor neuron (SMN) complex, which in turn facilitates spliceosome assembly and is associated with spinal muscular atrophy^43^. We derived three independent i^3^N iPSC clones containing bi-allelic deletions for this candidate PIR and observed significant downregulation of *STRAP* expression following differentiation of the i^3^N iPSCs into excitatory neurons (two sample t-test, two-sided, *p*=9.4×10^−13^). Targeting the same region with CRISPRi silencing also consistently downregulated *STRAP* expression in the excitatory neurons (two sample t-test, two-sided, *p*=1.5×10^−7^) (**Supplementary Fig. 5c**). Next, we focused on a candidate PIR 20 kb upstream from the promoter of *DRD2*, which encodes the D2 subtype of the dopamine receptor. Previously, rs2514218, a noncoding variant 47 kb upstream from *DRD2*, was found to be associated with anti-psychotic drug response in a cohort of schizophrenia patients^44^. Notably, this variant is in linkage with a cluster of SCZ SNPs overlapping open chromatin peaks within our candidate PIR. *DRD2* is also the gene associated with the Taq1A polymorphism which is linked to reduced dopamine receptor density as well as addiction, anxiety, depression, and social problems in patients^45^. We first demonstrate that mono-allelic deletion of the candidate PIR in three independent clones leads to significant downregulation of *DRD2* expression in excitatory neurons (two sample t-test, two-sided, *p*=2.7×10^−7^) (**Fig. 5e**). Next, we confirm that mono-allelic deletion of the candidate PIR leads to allelic imbalance in the expression of *DRD2* by performing TOPO cloning and genotyping cDNA with allele-specific variants (**Supplementary Fig. 5d**). *DRD2* is a key gene possessing multifaceted roles in human brain function, and it has been implicated in numerous complex neurological disorders including addiction, bipolar disorder, migraine, and obesity^46^. By prioritizing and validating putative regulatory SNPs for genes such as *DRD2*, our integrative approach enables the development of novel therapeutic and diagnostic strategies targeting specific variants for their roles in otherwise recalcitrant complex neurological disorders.

### Genetic variants contribute to chromatin interaction bias and alterations in gene expression

Since regulatory variants and other genetic perturbations are thought to disrupt chromatin looping between promoters and PIRs, we were interested to see if we could detect instances of allelic bias across our sets of significant promoter-PIR interactions. We used our chromatin interaction data to perform genome-wide phasing of WTC11 variants using HaploSeq^47^ and performed allele-specific mapping at a resolution of 10 kb using HiC-Pro. We identify 41 (0.185%) and 151 (0.703%) of significantly interacting bins to exhibit allelic bias at an FDR cutoff of 5% in excitatory neurons and lower motor neurons, respectively, confirming that genetic diversity can contribute to allelic bias in chromatin interactions (**Fig. 6a, Supplementary Table 10**). In one instance, allelically biased interactions were detected between a PIR containing SNPs for bipolar alcoholism^48^ and the promoter of *SYT17* (**Fig. 6b**). *SYT17* encodes a member of a family of membrane-trafficking proteins that mediate synaptic function and regulate calcium-controlled neurotransmitter release^49^. The identification of chromatin interactions with allelic bias at the *SYT17* locus suggests that variants can increase individual risk for bipolar alcoholism by disrupting interactions for *SYT17*, consistent with a model in which regulatory sequences recruit TFs and other activating factors to form physical contacts with their gene targets.

**Figure 6.**
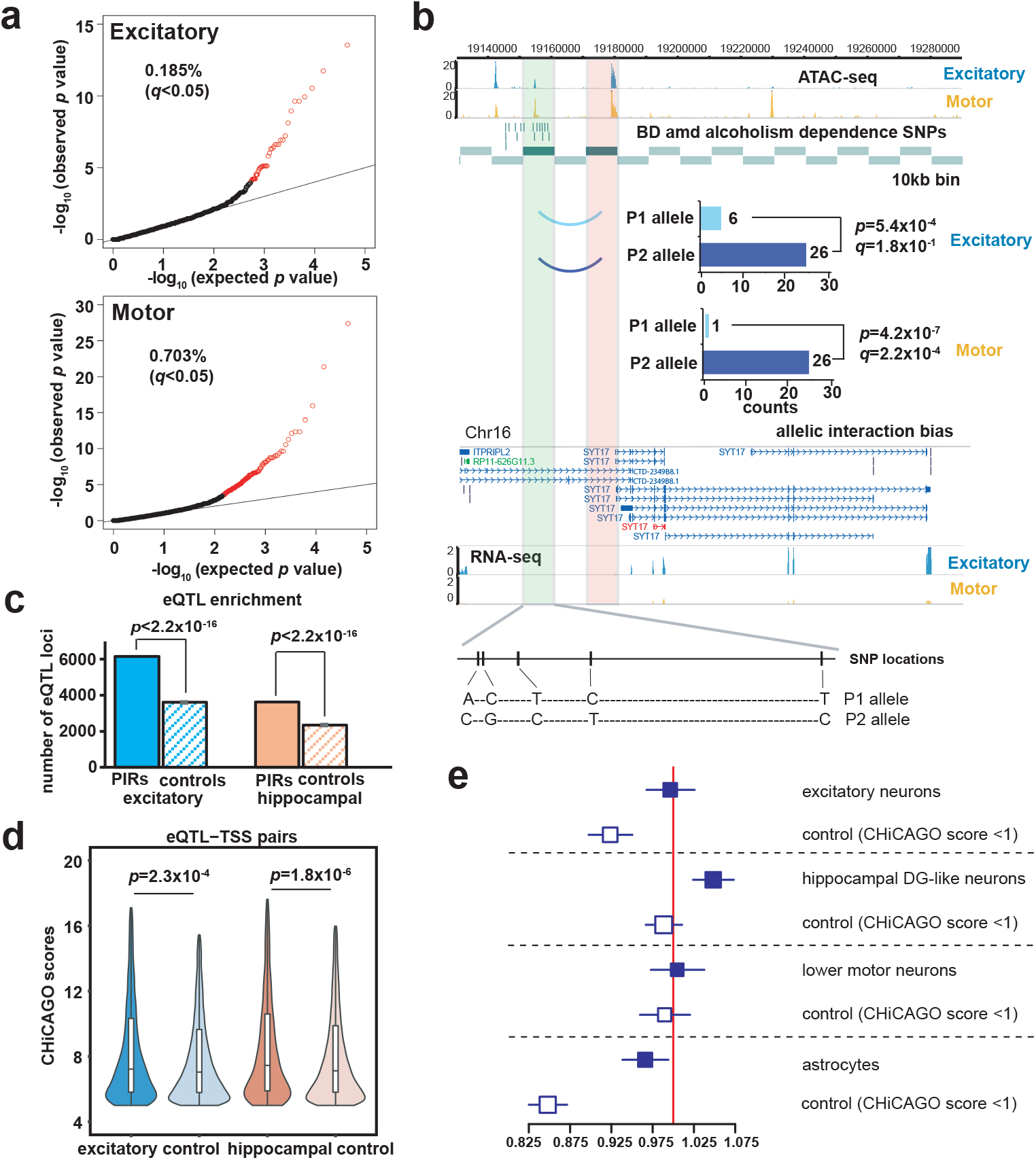
Genetics variants contribute to chromatin interaction bias and alterations in gene expression. (**a**) Quantile-quantile plots showing the proportions of interacting 10 kb bins exhibiting significant allelic bias at an FDR of 5% in excitatory neurons and lower motor neurons. (**b**) An example of an interaction exhibiting significant allelic bias in excitatory neurons (binomial test, two-sided, *p*=5.4×10^−4^) and lower motor neurons (binomial test, two-sided, *p*=4.2×10^−7^). The interaction occurs between a PIR containing SNPs for bipolar alcoholism at an open chromatin peak (green highlight) and the promoter of *SYT17* (orange highlight). Heterozygous WTC11 variants at the PIR are shown, along with bar graphs of detected read counts for each allele in our chromatin interactions. (**c**) Enrichment of significant eQTLs from GTEX V7 at significant versus randomly shuffled PIRs in matched tissue types for excitatory neurons and hippocampal DG-like neurons (one sample z-test, *p*<2.2×10^−16^ for both cell types). Error bars show the standard deviation over 100 sampled sets of randomly shuffled PIRs. (**d**) Distributions of interaction scores for chromatin interactions overlapping significant eQTL-TSS pairs versus randomly sampled nonsignificant eQTL-TSS pairs in excitatory neurons and hippocampal DG-like neurons. Interaction scores are significantly enriched for significant eQTL-TSS pairs (Kolmogorov-Smirnov test, *p*=2.3×10^−4^ for excitatory neurons and *p*=1.8×10^−6^ for hippocampal DG-like neurons). (**e**) Forest plot for independent ASD case-specific and non-overlapping matched pseudo-control-specific SNP pairs for each cell type. The x-axis shows the odds ratio (OR) estimated between the numbers of case- and control-specific SNP pairs at a significance threshold of 10^−7^ in each cell type. The area inside the squares is proportional to the number of observations for each comparison. Hippocampal DG-like neuron promoter-PIR pairs showed case-specific enrichments of ASD SNP pairs (chi-squared test, *p*<2.7×10^−16^). The confidence intervals for each OR estimation are shown in blue, and the red line represents a baseline OR of 1.

Physical chromatin interactions have also been theorized to mediate the effects of *cis*-acting regulatory variants such as expression quantitative trait loci (eQTLs) on gene expression. To test this hypothesis, we first show that significant eQTLs filtered at an FDR cutoff of 5% in cortical and hippocampal tissues from GTEx V7^50^ are enriched at PIRs for the excitatory neurons and hippocampal DG-like neurons, respectively (one sample z-test, *p*<2.2×10^−16^ for both cell types) (**Fig. 6c**). Next, we demonstrate that the mean scores for interactions overlapping significant eQTL-TSS pairs are significantly higher than the scores for interactions overlapping randomly shuffled eQTL-TSS pairs (Kolmogorov-Smirnov test, *p*=2.28×10^−4^ for excitatory neurons and *p*=1.76×10^−6^ for hippocampal DG-like neurons) (**Fig. 6d**). This indicates that significant promoter-PIR interactions recapitulating regulatory relationships between eQTL-TSS pairs are called with increased levels of confidence. Our results present orthogonal lines of evidence that chromatin interactions can not only be altered by variants in an allele-specific manner, but that variants can also influence gene expression through the formation of chromatin interactions.

### Enrichment of chromatin interactions at ASD SNP pairs

Genetic epistasis refers to combinations of independent variants exhibiting effects which cannot be predicted from their individual contributions alone, and it has been found to contribute to complex traits, including behavioral ones, in model organisms^51–54^. Epistasis has also been theorized to drive complex traits in humans such as ASD^55^. However, studying epistasis in humans in an unbiased fashion is challenging due to the large number of loci that need to be tested. In addition, most candidate pairs studied for epistasis are based on interactions between proteins encoded at the loci of interest. Therefore, chromatin interactions present a unique opportunity to identify epistatic effects in an unbiased fashion, as they suggest an attractive model in which physical contacts bridge independent variants to mediate genetic interactions in epistasis.

For each cell type, we tested for epistatic effects between all combinations of SNP pairs occurring at both ends of our significant promoter-PIR interactions. Specifically, we evaluated enrichment in our interactions for independent ASD case-specific SNP pairs over non-overlapping matched pseudo-control-specific SNP pairs (see methods)^56^. Of all the cell types, only the hippocampal DG-like neurons showed enrichment at PLINK’s default significance threshold of 10^−7^ (chi-squared test, *p*=2.7×10^−16^) (**Fig. 6e**). Next, we explored sets of increasingly stringent significance thresholds (**Supplementary Table 11**), demonstrating that excitatory neurons also exhibit enrichment at a significance threshold of 10^−20^ (chi-squared test, *p*=0.02). As an additional set of controls, we tested for epistatic effects in nonsignificant promoter-PIR interactions with score < 1, demonstrating that for all cell types, significant interactions exhibit greater enrichments for epistatic effects than are observed in nonsignificant ones. Overall, our results confirm that genetic interactions can indeed be mediated by chromatin interactions, underscoring the importance of the 3D epigenome for elucidating factors underlying individual disease risk.

## Discussion

There is a distinct lack of 3D epigenomic annotations in cell types that are relevant to disease and development, especially in the field of brain research. Past studies have frequently relied on heterogeneous tissues containing cell types with disparate functions, limiting their abilities to detect and interpret instances of cell type-specific gene regulation. Neurons and glia, for example, represent lineages with divergent functions that coexist in most tissues of the CNS. At the same time, complex diseases often involve multiple dysregulated loci exhibiting cell type-specific patterns of activity. This presents unique challenges for deciphering disease etiology, for example in distinguishing causative mechanisms from secondary, reactive phenotypes across distinct populations of cells. Therefore, the annotation of regulatory relationships in specific, well-characterized cell types should enable the wider community to derive numerous insights into complex disease biology. Chromatin interactions in particular are ideal for mapping promoters to distal regulatory elements, as they provide direct physical evidence of regulatory sequences contacting genes. To date, several studies have characterized chromatin interactions in fetal brain tissues and neural cell types^6,57^. However, these studies used *in situ* Hi-C, which lacks power for calling interactions compared to targeted approaches such as pcHi-C.

In this study, we leverage pcHi-C, ATAC-seq, and RNA-seq to annotate previously uncharted regulatory relationships between promoters and distal regulatory elements in cell types that are relevant to neurological disease. We show that PIRs in our interactions are not only cell type-specific, but they are also enriched for regulatory chromatin signatures including open chromatin peaks as well as *in vivo* validated enhancers in the VISTA Enhancer Browser. Despite functional similarities between the cell types used in this study, the inspection of cell type-specific distal open chromatin peaks at PIRs reveals subtype-specific binding sites for TFs involved in the specification and maintenance of cellular identity. Furthermore, our interactions enable the identification of novel gene targets for disease-associated variants and the prioritization of variants for validation using CRISPR techniques. We report a large number of putative regulatory variants which may reveal additional aspects of complex disease biology. Finally, the disease- and cell type-specific enrichment of variants at PIRs, combined with the observation that the same PIRs can target different promoters in different cell types, affirms that regulatory variants possess context-dependent functional specificities, underscoring the importance of performing validation in the appropriate cell types.

The integrative analysis in this study has several limitations including a lack of cell type-specific annotations for various genomic and epigenomic features occurring at PIRs. For example, the analysis of chromatin state enrichment at PIRs utilized data in matched tissues from the Roadmap Epigenomics Project. Although we generated our own maps of H3K27ac and CTCF binding sites, our results would be even more sensitive if chromatin states were inferred in matching cell types. Furthermore, while studying chromatin interactions in healthy cells enables the detection of regulatory interactions in the absence of dysregulation, the characterization of patient-derived iPSCs will also be important in the future to glean specific insights into how the 3D epigenome is altered in disease. Additional follow-up experiments are necessary to determine how the haploinsufficiency of proteins such as STRAP and DRD2 may contribute to phenotypes in disease. Finally, *in vitro* cultured cells can at present only approximate the full set of cellular responses occurring *in vivo*, especially in complex structures such as the brain. Future approaches isolating specific cell populations from tissues, leveraging single cell sequencing, or studying advanced organoid models will be essential for drilling down even deeper into mechanisms driving cellular identity and disease. In conclusion, our study presents a roadmap for the annotation of cell type-specific interactions in the CNS, advancing our ability to elucidate mechanisms by which noncoding variants drive complex neurological disorders. The epigenomic characterization of additional cell types should continue to yield rich insights into the landscape of transcriptional regulation, contributing towards an improved understanding of complex disease biology^58^.

## Methods

### Cell culture

Human excitatory neurons were generated using integrated, isogenic, and inducible neurogenin-2 (Ngn2) iPSCs (i^3^N iPSCs) with doxycycline-inducible mouse Ngn2 integrated at the AAVS1 safe-harbor locus. The i^3^N iPSCs have a well-characterized wild type genetic background (WTC11)^7^. A simplified, two-step pre-differentiation and maturation protocol was used to generate excitatory neurons^8^. Briefly, i^3^N iPSCs were incubated with 2 μg/ml doxycycline in pre-differentiation media containing KnockOut DMEM/F12 supplemented with 1x N-2, 1x NEAA, 1 μg/ml mouse laminin, 10 ng/ml BDNF, and 10 ng/ml NT3. In addition, 10 μM Rock inhibitor was included in the pre-differentiation media for the first day. Media was changed daily for three days. For maturation, pre-differentiated precursor cells were dissociated and subplated on poly-D-lysine/laminin plates in maturation media containing equal parts DMEM/F12 and Neurobasal-A with 2 μg/ml doxycycline and supplemented with 0.5x B-27, 0.5x N-2, 1x NEAA, 0.5x GlutaMax, 1 μg/ml mouse laminin, 10 ng/ml BDNF, and 10 ng/ml NT3. The doxycycline was omitted from all subsequent media changes. Half of the media was half changed weekly over the first two weeks, then the amount of media was doubled on day 21. Thereafter, a third of the media was replaced weekly until harvesting. 7 to 8 week old excitatory neurons were harvested for library preparation.

Human hippocampal DG-like neurons were generated from dissociated hippocampal organoids (unpublished). Briefly, WTC11 iPSCs were grown on MEF feeder cells and patterned towards a neural ectoderm fate using dual SMAD inhibition as floating embryoid bodies (EBs) in media containing 20% KnockOut Serum Replacement. Next, 4 week old EBs were patterned towards a hippocampal fate using WNT and BMP in media containing 1x N-2. After patterning, organoids were dissociated using a neural tissue dissociation kit (MiltenyiBiotech), plated on PDL- and laminin-coated plates, then cultured for 4 weeks in media containing 1x B-27, 10 ng/ml BDNF, 10 ng/ml GDNF, 0.5 mM cAMP, and 200 μM ascorbic acid.

Human lower motor neurons were differentiated from WTC11 iPSCs with a doxycycline inducible transgene expressing NGN2, ISL1, and LHX3 integrated at the AAVS1 safe-harbor locus (i^3^LMN iPSCs) as previously reported^10^. Briefly, i^3^LMN iPSCs were maintained on growth factor reduced Matrigel in StemFit media (Nacalai USA). On day 0, 1.5×10^6^ i^3^LMN iPSCs were plated on 10-cm dishes, followed 24 hours later by exchange into neural induction media containing doxycycline and compound E. On day 3, the precursor cells were replated onto 12-well plates coated with poly-D-lysine and laminin at a density of 2.5×10^5^ cells per well. From day 3 to day 4, the cells were treated with a pulse of 40 μM BrdU for 24 hours to suppress the proliferation of undifferentiated cells. Media was exchanged on day 4 and every three days thereafter. The cells were harvested 10 days post-differentiation for library preparation.

Human primary astrocytes (P0) were purchased from ScienCell Research Laboratories (catalog #1800) and cultured using the recommended media (catalog #1801). Briefly, cells were cultured in flasks coated with poly-L-lysine (2μ/cm^2^) and passaged once using trypsin and EDTA before harvesting.

All cells used in the present study were verified as mycoplasma contamination free.

### Immunofluorescence

Cells were fixed in 4% paraformaldehyde (PFA) for 15 minutes at room temperature, then washed multiple times with PBS containing 0.1% Triton X-100 (PBS-T) before undergoing blocking using a solution of 5% bovine serum albumin (BSA) in PBS-T for 1 hour at room temperature. Primary antibodies against Cux1 (Abcam, ab54583, lot: GR3224721-2), MAP2 (Abcam, ab5392, lot: GR3242762-1), PROX1 (Millipore, MAB5654, lot: 3075604), HB9 (Millipore, ABN174, lot: 3050643), SMI32 (Abcam, ab7795, lot: GR299862-23), and GFAP (Abcam, ab7260, lot: GR3240356-1) were diluted in 5% BSA solution and incubated overnight at 4°C prior to use. Secondary antibodies including Alexa Fluor 568 goat anti-chicken IgG, Alexa Fluor 568 goat anti-mouse IgG, Alexa Fluor 488 donkey anti-rabbit IgG, and Alexa Fluor 488 donkey anti-mouse IgG (Molecular Probes) were diluted in 5% BSA solution and incubated for 1 to 2 hours at room temperature prior to use. Images were acquired using a Leica TCS SP8 confocal microscope with a 40x oil immersion objective lens.

### Promoter capture Hi-C (pcHi-C)

*In situ* Hi-C libraries for the excitatory neurons, hippocampal DG-like neurons, lower motor neurons, and astrocytes were constructed from 1 to 2 million cells using HindIII as a restriction enzyme as previously described^59^. pcHi-C was performed using biotinylated RNA probes prepared according to an established protocol (Jung et al., under review). Briefly, sets of 120 bp probes with 30 bp overhangs were designed to capture the sequences adjacent to restriction sites flanking each promoter-containing HindIII fragment. Three probes were targeted to each side of each restriction site, such that a total of 12 probes targeted each promoter-containing HindIII fragment. In total, promoters (defined as the sequences up to 500 bp upstream and downstream of each transcription start site) for 19,603 of the 20,332 protein coding genes in GENCODE 19 were captured using this approach. While noncoding RNA promoters were not explicitly targeted by this design, HindIII fragments containing 3,311 of the 14,069 noncoding RNA promoters in GENCODE 19 were also baited by the probes.

To perform the hybridization, 500 ng of *in situ* Hi-C libraries were first mixed with 2.5 μg human Cot-1 DNA (Invitrogen #15279011), 2.5 μg salmon sperm DNA (Invitrogen #15632011), and 0.5 nmol each of the p5 and p7 IDT xGen Universal Blocking Oligos in a volume of 10 μL, then denatured for 5 min at 95°C before holding at 65°C. Next, a hybridization buffer mix was prepared by mixing 25 μL 20x SSPE, 1 μL 0.5 M EDTA, 10 μL 50x Denhardt’s solution, and 13 μL 1% SDS, followed by pre-warming to 65°C. Finally, 500 ng of the probe mix was combined with 1 μL 20 U/μL SUPERase-In (Invitrogen #AM2696) in a 6 μL volume, pre-warmed to 65°C, then promptly mixed with the library and hybridization buffer mix. The final solution was transferred to a humidified hybridization chamber and incubated for 24 hours at 65°C. 0.5 mg Dynabeads MyOne Streptavidin T1 magnetic beads (Invitrogen #65601) were used to pull down the captured fragments in a binding buffer consisting of 10 mM Tris-HCl pH 7.5, 1 M NaCl, and 1 mM EDTA. Next, the beads were washed once with 1x SSC and 0.1% SDS for 30 minutes at 25°C, followed by three washes with pre-warmed 0.1X SSC and 0.1% SDS for 10 minutes each at 65°C. The final library was eluted in 20 μL nuclease-free water, amplified, then sent for paired-end sequencing on the HiSeq 4000 (50 bp reads), the HiSeq X Ten (150 bp reads), or the NovaSeq 6000 (150 bp reads).

A detailed description of the capture probe design and experimental procedures can be viewed in the attached manuscript which is under review in Nature.

### ATAC-seq

ATAC-seq was carried out as previously described using the Nextera DNA Library Prep Kit (Illumina #FC-121-1030)^60^. First, frozen or fresh cells were washed once with ice cold PBS containing 1x protease inhibitor before being exchanged into ice cold nuclei extraction buffer (10 mM Tris-HCl pH 7.5, 10 mM NaCl, 3 mM MgCl_2_, 0.1% Igepal CA630, and 1x protease inhibitor) and incubated for 5 minutes on ice. Next, 50,000 cells were counted out, exchanged into 1x Buffer TD, then incubated with 2.5 μL TDE1 for 30 minutes at 37°C with shaking. The transposed DNA was purified using Qiagen MinElute spin columns, amplified with Nextera primers, then size-selected for fragments between 300 and 1000 bp in size using AMPure XP beads. Libraries were sent for single-end sequencing on the HiSeq 4000 (50 bp reads) or paired-end sequencing on the NovaSeq 6000 (150 bp reads). Sequencing reads were mapped to hg19 and processed using the ENCODE pipeline (https://github.com/kundajelab/atac_dnase_pipelines) running the default settings. Only the first read was used, and all sequencing reads were trimmed to 50 bp prior to mapping. Open chromatin peaks called by the pipeline were expanded to a minimum width of 500 bp for all downstream analyses. Peaks overlapping coding gene or noncoding RNA promoters were assigned as promoter open chromatin peaks, while all other peaks were assigned as distal open chromatin peaks. All data processing metrics are reported in **Supplementary Table 1**.

### RNA-seq

RNA was extracted using the RNeasy Mini Kit (Qiagen #74104). Approximately 500 ng of extracted RNA was used to construct libraries for sequencing using the TruSeq Stranded mRNA Library Prep Kit (Illumina #20020594). Libraries were sent for single-end sequencing on the HiSeq 4000 (50 bp reads) or paired-end sequencing on the NovaSeq 6000 (150 bp reads). Raw sequencing reads were aligned to hg19/GRCh37 using STAR running the standard ENCODE parameters, and transcript quantification was performed in a strand-specific manner using RSEM with the annotation from GENCODE 19. Only the first read was used, and all sequencing reads were trimmed using TrimGalore 0.4.5 running the following options: -q 20 --length 20 --stringency 3 --trim-n. The *edgeR* package in R was used to calculate TMM-normalized RPKM values for each gene based on the expected counts and gene lengths for each replicate reported by RSEM. The mean gene expression across all replicates was used for each cell type. All data processing metrics are reported in **Supplementary Table 1**.

### CUT&RUN

CUT&RUN libraries for excitatory neurons and lower motor neurons were constructed for 100,000 to 250,000 cells with antibodies for H3K27ac and CTCF as previously described^24^. First, cells were lysed in nuclei extraction buffer (20 mM HEPES-KOH pH 7.9, 10 mM KCl, 1 mM MgCl_2_, 0.1% Triton X-100, 20% glycerol, and 1x protease inhibitor) for 10 minutes on ice. Next, samples were spun down and washed twice with nuclei extraction buffer before being resuspended in 100 μL nuclei extraction buffer. 10 μL of Concanavalin A-coated beads previously washed and resuspended in binding buffer (1x PBS, 1 mM CaCl2, 1 mM MgCl2, and 1 mM MnCl2) were then added to the samples and incubated with rotation for 15 min at 4°C. Next, samples were washed once each with Buffer 1 (20 mM HEPES-KOH pH 7.9, 150 mM NaCl, 2 mM EDTA, 0.5 mM spermidine, 0.1% BSA, and 1x protease inhibitor) and Buffer 2 (20 mM HEPES-KOH pH 7.9, 150 mM NaCl, 0.5 mM spermidine, 0.1% BSA, and 1x protease inhibitor) before being resuspended in 50 μL of Buffer 2 containing 0.5 μL antibody (H3K27ac from Active Motif, 39122, lot: 22618011 and CTCF from Millipore 07-729, lot: 305960) and incubating for at least 2 hours with rotation at 4°C. Following the incubation, samples were washed twice with Buffer 2 before being incubated in 50 μL of Buffer 2 containing ~700 ng/mL protein A-MNase fusion protein (Batch #6 from the Henikoff Lab) for 1 hour with rotation at 4°C. Samples were washed two more times and resuspended in 100 μL of Buffer 2 before starting the MNase digestion by adding CaCl2 to a concentration of 2 mM (with the samples kept on ice), followed 30 minutes thereafter by the addition of 100 μL 2X Stop Buffer (200 mM NaCl, 20 mM EDTA, 4 mM EGTA, 50 ug/mL RNase A, 40 ug/mL glycogen, and 2 pg/mL spike-in DNA) to inactivate the MNase. Samples were incubated for 20 min at 37°C and spun down for 5 minutes at 4°C to release DNA fragments that were subsequently extracted from the supernatant using Qiagen MiniElute spin columns. Libraries were prepared using TruSeq adapters and size-selected using SPRIselect beads before being amplified and sent for paired-end sequencing on the NovaSeq 6000 (150 bp reads). Sequencing reads were first trimmed to 50 bp using fastp then mapped to hg19 using bowtie2 running the following options: --local --very-sensitive--local --no-mixed --no-discordant -I 10 -X 700. Picard Tools was used to remove duplicate reads, and MACS2 was used to call peaks on merged replicates at an FDR cutoff of 5%.

### Validation of PIRs using CRISPR deletion

To validate genomic interactions captured by pcHi-C, candidate PIRs were targeted for CRISPR deletion in i^3^N iPSCs. At each locus of interest, we designed pairs of sgRNAs to delete the putative regulatory element as localized by open chromatin peaks in the candidate PIR. All sgRNAs were synthesized by Synthego, and Cas9 protein was sourced from QB3-Berkeley. To generate deletion lines, CRISPR/Cas9 nucleofections were performed using the LONZA Human Stem Cell Nucleofector^®^ Kit. For each nucleofection, 500,000 i^3^N iPSCs were transfected with Cas9:sgRNA RNP complex (consisting of 12 μg Cas9, 10 μg sgRNA 1, and 10 μg sgRNA 2) using program “A-023” on the LONZA 4D-Nucleofector. The cells were then seeded onto Matrigel-coated 6-well plates containing Essential 8™ Medium (ThermoFisher #A15169-01) with added Y-27632 for recovery following nucleofection. After 48 hours, the cells were split into new 6-well plates at a concentration of approximately 50 cells per well for picking single colonies. Clones picked from the 6-well plates containing homozygous deletions were confirmed by PCR and induced into excitatory neurons for quantifying the expression of genes targeted by the deleted elements. We used three independent deletion clones for each experiment, and clones with wild type genotypes were used as controls. To perform the quantification, total RNA from the excitatory neurons was extracted using a Qiagen AllPrep DNA/RNA Mini Kit, and cDNA was synthesized using a Bio-RAD iScript™ cDNA Synthesis Kit. qPCR for targeted genes was performed with FastStart Essential DNA Green Master reaction mix (Roche) on the LightCycler^®^ 96 System (Roche). All CRISPR deletion experiments were performed with two independent transfections. Detailed information on all the primers used can be found in **Supplementary Table 12**.

### Validation of PIRs using CRISPRi

Excitatory neurons induced from i^3^N iPSCs were infected with lentivirus carrying dCas9-KRAB-blast (Addgene #89567), and colonies with high expression of dCas9 were picked. The CROP-seq-opti vector (Addgene #106280) was used for sgRNA expression. sgRNAs were designed, cloned, and cotransfected with lentivirus packaging plasmids pMD2.G (Addgene #12259) and psPAX (Addgene #12260) into 293T cells using PolyJet (SignaGen Laboratories #SL100688) according to the manufacturer’s instructions. Virus-containing media was collected for 72 hours, filtered through a 0.45 μm filter (Millipore #SLHV033RS), and concentrated using an Amicon Ultra centrifugal filter (Millipore #UFC801024). The virus was titrated into the excitatory neurons by qPCR 72 hours after infection. The internal control for qPCR targeted an intronic region (forward primer: TCCTCCGGAGTTATTCTTGGCA, reverse primer: CCCCCCATCTGATCTGTTTCAC). Integration of the WPRE fragment was quantified in comparison with a control cell line containing a known copy number of WPRE. For CRISPRi silencing of putative regulatory elements, excitatory neurons were treated with lentivirus containing sgRNAs (MOI ~3). Cells were collected for mRNA extraction 7 days following transfection, and the expression of target genes was determined by qPCR. All CRISPRi experiments were performed in triplicate, with three technical replicates per experiment. Detailed information on all the primers used can be found in **Supplementary Table 12**.

### Reproducibility and saturation analysis

We took pcHi-C contact matrices generated at 10 kb resolution using HiC-Pro 2.11.0 with the following settings (MIN_MAPQ=20, MIN_FRAG_SIZE=100, MAX_FRAG_SIZE=5000000, MIN_INSERT_SIZE=100, MAX_INSERT_SIZE=1200, and reporting only bin pairs that are baited on at least one end with our pcHi-C probes, with all other settings set to their default values) and calculated the pairwise stratum adjusted correlation coefficient (SCC) between replicates across all cell types using HiCRep 1.4.0 on chromosome 1 (h=20 and only considering contacts with distances below 5 Mb). SCCs evaluated on the other chromosomes closely resembled the results for chromosome 1 (data not shown). Hierarchical clustering for the pairwise SCC values was performed using the Seaborn *clustermap* function in Python. Pairwise correlation heatmaps and clustering dendrograms for ATAC-seq replicates were generated by counting reads overlapping a set of consensus peaks using the *DiffBind* package in R, with the set of consensus peaks defined as peaks occurring in at least two replicates across all cell types (minOverlap=2). Pairwise distance estimates and clustering dendrograms for RNA-seq replicates were generated using the *DESeq2* package in R. For saturation analysis, we first downsampled all pcHi-C libraries to 5%, 10%, 20%, 40%, 60%, 80%, and 100% of the final sequencing depths used in the study. Next, we computed pairwise SCCs between all pairs of biological replicates using HiCRep at these downsampled sequencing depths.

### Calling significant promoter-PIR interactions

Paired-end sequencing reads were first trimmed using fastp running the default settings before being mapped, filtered, and deduplicated using HiCUP v0.71 with bowtie2 and filtering for ditags between 100 and 1200 bp^61^. In addition, the sequencing depths of all libraries was normalized so that each replicate had the same number of usable reads, or the number of on-target cis pairs interacting over a distance of 10 kb. Significant promoter-PIR interactions were called using CHiCAGO running the default settings, retaining baited fragments that are supported by at least 250 reads (minNPerBaits=250). Promoter-PIR interactions between HindIII fragments with a score (negative log p-value) of 5 or greater in each cell type were determined to be significant. All data processing metrics are reported in **Supplementary Table 1**. In cases where CHiCAGO reported the same interaction twice due to directionality between two bait-containing fragments (i.e. bait A to bait B, bait B to bait A), the two interactions were merged, retaining the more significant score of the two interactions. Interchromosomal interactions were also omitted from the analysis. To call overlaps between our sets of significant interactions and genomic and epigenomic features including promoters, open chromatin peaks, chromatin states, disease-associated variants, and eQTLs, interacting bins were expanded to a minimum width of 5 kb or retained as the original widths of the HindIII fragments if they exceeded 5 kb. Interactions involving HindIII fragments larger than 100 kb were omitted from our analysis due to low resolution. An interaction was considered to be shared between cell types if both its interacting ends intersected the corresponding ends of an interaction in another cell type. Otherwise, an interaction was classified to be cell type-specific.

### Chromatin state analysis

Annotations for the publicly available 15 state ChromHMM model were downloaded from the Roadmap Epigenomics Project for the dorsolateral prefrontal cortex (E073, “Brain Dorsolateral Prefrontal Cortex”), hippocampus (E071, “Brain Hippocampus Middle”), and normal human astrocytes (E125, “NH-A Astrocytes Primary Cells”). The states were available at a resolution of 200 bp and grouped as follows: TssA and TssAFlnk were merged as TSS, TxFlnk, Tx, and TxWk were merged as Tx, EnhG and EnhBiv were merged as other enhancer, and ReprPC and ReprPCWk were merged as ReprPC. All other states (Enh, ZNF/Rpts, Het, TssBiv, and BivFlnk) were used as is. Enrichment analysis was performed for each cell type by counting the number of chromatin states overlapping significant PIRs versus the number of chromatin states overlapping randomly shuffled PIRs with matching distance distributions. A total of 100 sets of randomly shuffled PIRs were sampled in each case.

### GO enrichment analysis

Protein coding and noncoding RNA genes from GENCODE 19 participating in significant cell type-specific promoter-PIR interactions were used for cell type-specific GO enrichment analysis. Only genes participating in interactions between promoter-containing and non-promoter-containing bins with a promoter open chromatin peak on one end and a distal open chromatin peak on the other end were used. The promoter open chromatin peaks were used to define the genes with promoters interacting with cell type-specific PIRs. A minimum normalized RPKM of 0.5 was used to filter out genes not significantly expressed in each cell type, and the resulting gene lists were input into Enrichr. Enriched GO terms from the “GO Biological Process 2018” ontology are reported according to their combined scores (calculated by multiplying the log of the p-value by the z-score of the deviation from the expected rank). For disease-specific GO enrichment analysis, target genes across all cell types were combined and input into Enrichr. All promoters in GENCODE 19 were included in this analysis. The top 100 enriched cell type-specific and disease-specific GO terms for each category and their raw p-values are reported in **Supplementary Table 3** and **Supplementary Table 9**.

### Motif enrichment analysis

We took the sets of all cell type-specific distal open chromatin peaks participating in significant promoter-PIR interactions between promoter-containing and non-promoter-containing bins for each cell type, and used the sequences in 250 bp windows around the peak summits to perform motif enrichment analysis using HOMER running the default settings. The entire genome was used as a background. Significance and expression values for each detected motif and its corresponding TFs are reported in **Supplementary Table 3**. Entries with similar or identical consensus TF motif sequences were grouped for brevity.

### VISTA enhancer analysis and target gene identification

Human enhancer regions and mouse enhancer regions with orthologous human sequences associated with positive annotations in the VISTA Enhancer Browser were downloaded and analyzed for overlap with our sets of significant promoter-PIR interactions for each cell type. Of the 2,956 total tested elements in their database (January 2019), 1,568 were found to be positive (976 were human elements and 892 were mouse elements with orthologous human sequences). Positive elements (expanded to a minimum width of 5 kb) found to participate in significant interactions are reported in **Supplementary Table 5**. For determining whether positive elements interacted with their nearest genes or with more distal genes, we only considered protein coding and noncoding RNA genes in GENCODE 19. To evaluate cases where interactions between positive elements and their nearest genes were unresolvable (“same fragment ambiguity”), we determined if a promoter for the nearest gene overlapped at least one HindIII fragment that the expanded positive element did not also overlap. The following were considered to be neural annotations: neural tube, hindbrain, cranial nerve, midbrain, forebrain, mesenchyme derived from neural crest, dorsal root ganglion, and trigeminal V.

### SNP enrichment analysis and target gene identification

GWAS SNPs for a total of eleven neurological disorders including Alzheimer’s disease (AD), attention deficit hyperactivity disorder (ADHD), amyotrophic lateral sclerosis (ALS), autism spectrum disorder (ASD), bipolar disorder (BD), epilepsy (EP), frontotemporal dementia (FTD), mental processing (MP), Parkinson’s disease (PD), and schizophrenia (SCZ), and unipolar depression (UD) were mined from the GWAS Catalog (December 2018) using a p-value threshold of 10^−6^. The GWAS SNPs were imputed using HaploReg v4.1 at an LD threshold of 0.8 according to the reported study population(s) for each SNP. The imputed SNPs were lifted over to hg19 and filtered for unique SNPs by position. See **Supplementary Table 5** for a detailed summary of the imputation process and the list of studies used. Disease- and cell type-specific enrichment for SNPs was calculated by taking the ratio of the number of SNPs overlapping significant PIRs over the mean number of SNPs over the number of SNPs overlapping randomly shuffled PIRs with matching distance distributions. A total of 100 sets of randomly shuffled PIRs were sampled in each case. To determine whether a GWAS SNP potentially interacted with a target gene, we determined whether it or any of its linked SNPs (expanded to a minimum width of 1 kb) interacted with a promoter for the nearest gene. To evaluate cases where interactions between GWAS SNPs and their nearest genes were unresolvable (“same fragment ambiguity”), we determined if a promoter for the nearest gene overlapped at least one HindIII fragment that a GWAS SNP or any of its linked SNPs did not also overlap. Finally, we derived a list of SNPs for which the SNP was located within 2 kb of the center of an open chromatin peak at a PIR, indicating strengthened evidence for a functional regulatory variant at that locus (“putative regulatory SNPs”).

### Phasing of the WTC11 genome

The raw WTC11 genome sequence can be downloaded from http://genome.ucsc.edu/cgi-bin/hgTracks?db=hg38&hubClear=https://s3-us-west-2.amazonaws.com/downloads.allencell.org/genome-sequence/ucsc_hubs/WTC_genome_hub/hub.txt. Phasing of the WTC11 genome was performed as previously described^47^. Briefly, WTC11 variants were first split by chromosome, and phase-informative pcHi-C reads were extracted using extractHAIRS with the minimum mapping quality set to 10 and the maximum insert size set to 30000000 bp^62^. Phasing was performed using Hapcut with a maximum of 101 iterations. Next, we extracted the maximum variants phased (MVP) haplotype block from the output of Hapcut to use as the seed haplotype. We modified the “neighborhood correction” aspect of phasing by filtering phased variants whose predicted phase would have a marginal probability below 0.99 using an in-house implementation of a hidden Markov model (HMM) as described previously^63,64^ with a reference haplotype set from the 1000 Genomes Project. Finally, missing variants were imputed using the same HMM with the reference haplotype set from the 1000 Genomes Project. The chromosome-wide SNP phasing data is available at the Gene Expression Omnibus under the following accessible number: GSE113483.

### Allelic bias analysis

We utilized the phasing data for the WTC11 genome along with the allele-specific mapping capabilities of HiC-Pro to quantify genome-wide allelic bias between significantly interacting 10 kb bins in excitatory neurons and lower motor neurons. We selected these two cell types because they used homogenous induction of TFs for differentiation, therefore minimizing the noise introduced by conventional differentiation techniques. Briefly, reads were mapped using bowtie to a version of the hg19 reference genome where all sites containing heterozygous phased SNPs were first N-masked. The unfiltered HiC-Pro contact maps were used for this analysis. Next, nucleotides at the masked polymorphic sites were used to assign the reads to either allele, with reads containing conflicting allele assignments or unexpected bases omitted from further analysis. Read pairs with at least one allele-specific mate were used to construct allele-specific Hi-C contact maps at 10 kb resolution, for which interacting bins overlapping with the set of significant promoter-PIR interactions with score ≥ 3 was used to detect bias. Only interacting bins with 10 or more reads across both alleles were kept. A two-sided binomial test was performed to assess allelic bias for each pair of interacting bins, and the resulting p-values were adjusted using the BH method to filter out significantly biased loci at an FDR cutoff of 5%. All allelically biased interactions with p-values < 0.001 are reported in **Supplementary Table 10**.

### eQTL enrichment analysis

1D enrichment of significant eQTLs with an FDR cutoff of 5% from GTEX V7 at significant versus randomly shuffled PIRs in matched tissue types for excitatory neurons (Brain - Cortex, n=136) and hippocampal DG-like neurons (Brain - Hippocampus, n=111) was performed similarly to the chromatin state and SNP enrichment analysis. Overall, we found that 4.7% of significant cortical eQTLs and 6.7% of hippocampal eQTLs interact in excitatory neurons and hippocampal DG-like neurons, respectively. To determine the 2D enrichment of eQTL-TSS pairs in our significant interaction sets, we first filtered out eQTL-TSS pairs that were within 10 kb of each other or on the same HindIII fragment as this would be below the minimum detectable resolution by pcHi-C. Next, we sampled a set of nonsignificant eQTL-TSS pairs with a matching distance distribution as the set of significant eQTL-TSS pairs for each cell type. We also controlled for the number of genes around which the eQTL-TSS pairs were centered. Finally, we compared the distributions of scores for significant interactions supporting the significant and nonsignificant sets of eQTL-TSS pairs by overlapping the eQTL-TSS pairs with our significant interactions.

### Epistasis analysis

In order to determine whether genetic variation in physically interacting regions might contribute to neurodevelopmental disorders via genetic interactions, we utilized GWAS data for ASD. For epistasis testing, we needed individual-level genotype data, so we used a dataset of 4,109 trios and 4,471,807 imputed and genotyped single nucleotide polymorphisms (SNPs), as previously reported^56^. This dataset includes publicly available ASD GWAS data [Autism Genetic Resource Exchange (AGRE), Autism Genome Project (AGP), Simons Simplex Collection (SSC)] in addition to in-house generated data [University of California, San Francisco (UCSF)], harmonized, imputed, and quality controlled (QC+) by us, as previously described^56^. These data comprised trios with one ASD-affected offspring and both parents selected for homogeneous genetic ancestry by multidimensional scaling with PLINK^65^. As our data were family-based and did not include unrelated controls, we used non-transmitted parental alleles, commonly known as pseudo-controls, generated using the *-tucc* option in PLINK. These 4,109 pseudo-controls are perfectly matched to ASD cases for ancestry, thereby serving as a control for any population confounding.

For each significant promoter-PIR interaction for each cell type, we extracted all independent SNPs in each region from our QC+ imputed GWAS data. We performed a case-only test for pairwise epistasis using the -*fast-epistasis, case-only*, and *set-by-set* options in PLINK with *p* ≤ 10^−7^ as a default threshold in ASD cases and in matched pseudo-controls^56^. Across the interacting loci in the four neural cell types, we performed approximately 19.7 million epistasis tests and 12,637,825 SNP pairs showed *p* ≤ 10^−7^ (65% tests performed).

Because all pairs of regions we tested were on the same chromosome (linked), we expected an excess of false positives (e.g. 65% at *p* ≤ 10^−7^) for the case-only test due to LD or haplotypes containing rare variants. We thus excluded all SNP pairs that showed epistasis with *p* ≤ 10^−7^ in both cases and pseudo-controls to generate case-specific and control-specific epistasis results for comparison, resulting in 0.5% of results with *p* ≤ 10^−7^. We next applied the –*clump* PLINK option to filter SNPs involved in potential epistasis in LD (r^2^ > 0.2) across pairs (e.g. one promoter SNP putatively interacting with several correlated PIR SNPs), such that only one pair (selected based on epistatic *p*-value) represented the combination of loci. This resulted in reduction of 15 to 19% of the number of pairs considered. The average distance between epistatic SNPs was 104 kb in excitatory neurons, 97 kb in hippocampal DG-like neurons, 87 kb in lower motor neurons, and 90 kb in astrocytes.

To test for enrichment of epistasis signal specific to ASD, we wanted to compare the number of SNP pairs at *P* ≤ 10^−7^ between the matched cases and pseudo-controls at various signal-to-noise ratios. We divided the set of case-specific epistatic SNP pairs and the set of control-specific epistatic SNP pairs into bins based on *P*-values [10^−30^ ≤ *p* ≤ 10^−7^] and a homogeneous number of SNP pairs to distinguish expected signal-to-noise (**Fig. 6e, Supplementary Table 7**). We then compared the number of case-specific and control-specific SNP pairs in each significance category using a proportion test in R. We performed meta-analysis of the case-specific and pseudo-control specific results across the four cell types using the *metafor* package in R. We illustrated the odds ratios of enrichment with the *forestplot* package in R.

As a negative control, we utilized equivalent promoter-PIR interactions with scores < 1 across the neural cell types. We sampled the same number of non-significant promoter-PIR pairs that we used in the main analysis for each cell type. For each non-significant interaction in each cell type, we filtered out those that overlapped interactions with score > 3 in any of the neural cell types. We also made sure to sample nonsignificant interactions with a similar interaction distance distribution as the significant interactions. We then performed SNP extraction, epistasis testing, and case-control enrichment analysis as described above for the nonsignificant interactions. Before filtering, 15-19% of SNPs showed *p* ≤ 10^−7^, and after filtering 7% met this threshold.

### Code availability statement

A copy of the custom code used for all the data analysis and figure generation in this study can be viewed and downloaded at the following GitHub repository: https://github.com/stayingsong/brain_pchic

### Data availability statement

All datasets used in this study (pcHi-C, ATAC-seq, RNA-seq, CUT&RUN, and chromosome-wide SNP phasing data) are available at the Gene Expression Omnibus under the accession number GSE113483. The reviewer access token is mjmrcsuaddkfhut.

Data can also be visualized on the WashU Epigenome Browser at the following link: http://epigenomegateway.wustl.edu/legacy/?genome=hg19&session=zEdB7v5de4&statusId=33592151

Tracks include ATAC-seq signal, RNA-seq plus/minus strand signal, CTCF CUT&RUN signal, and promoter-PIR interactions with score ≥ 5 for each cell type. HindIII fragments, positive Vista elements, GENCODE 19 genes, and SNPs for each disease are also shown.

## Acknowledgements

We thank Anthony Schmitt and Bing Ren for sharing pcHi-C probes and the pcHi-C protocol. Genomic analysis of the WTC11 line in this study was made possible by the whole genome sequencing data generated by the Allen Institute for Cell Science. We thank the Institute and its founder Paul G. Allen for making this work possible. We also thank G. Hon and S. Henikoff for providing reagents and Y. Guo, N. Ahituv, R. D. Hawkins, M. McManus, and B. Ren for providing critical feedback on the manuscript. This work was made possible in part by NIH-NEI EY002162 (Core Grant for Vision Research and the Research to Prevent Blindness Unrestricted Grant). This work was supported by the US National Institute of Health (NIH) grants R01AG057497 to YS, LG, and HS, R01EY027789 and UM1HG009402 to YS, the UCSF Weill Institute for Neuroscience Innovation Award, the Hillblom Foundation, and the American Federation for Aging Research New Investigator Award in Alzheimer’s Disease to YS. R01EY028249, R01HL130533, R01-HL13535801 to BRC as well as P01NS097206 and U19MH106434 to HS. R01MH105128, R35NS097370, and U19AI131130 to GLM. MS is supported by T32GM007175. FJ is supported by an NIH training grant T32GM73009.

## Author contribution

MS and YS designed the study. MS, XY, XR, LM, IJ, KJ and TWT performed the experiments. MS, BL, and IJ performed data analysis. LW supervised the epistasis analysis by MT. JD contributed to genomic phasing using HaploSeq. SL, JY, KW, BRC, FJ, GLM, HS, LD, CW, and LG contributed by providing cell samples. MS and YS prepared the manuscript with assistance from all authors.

## Competing interests

The authors declare no competing financial interests.

---

**Supplementary Figure 1.**
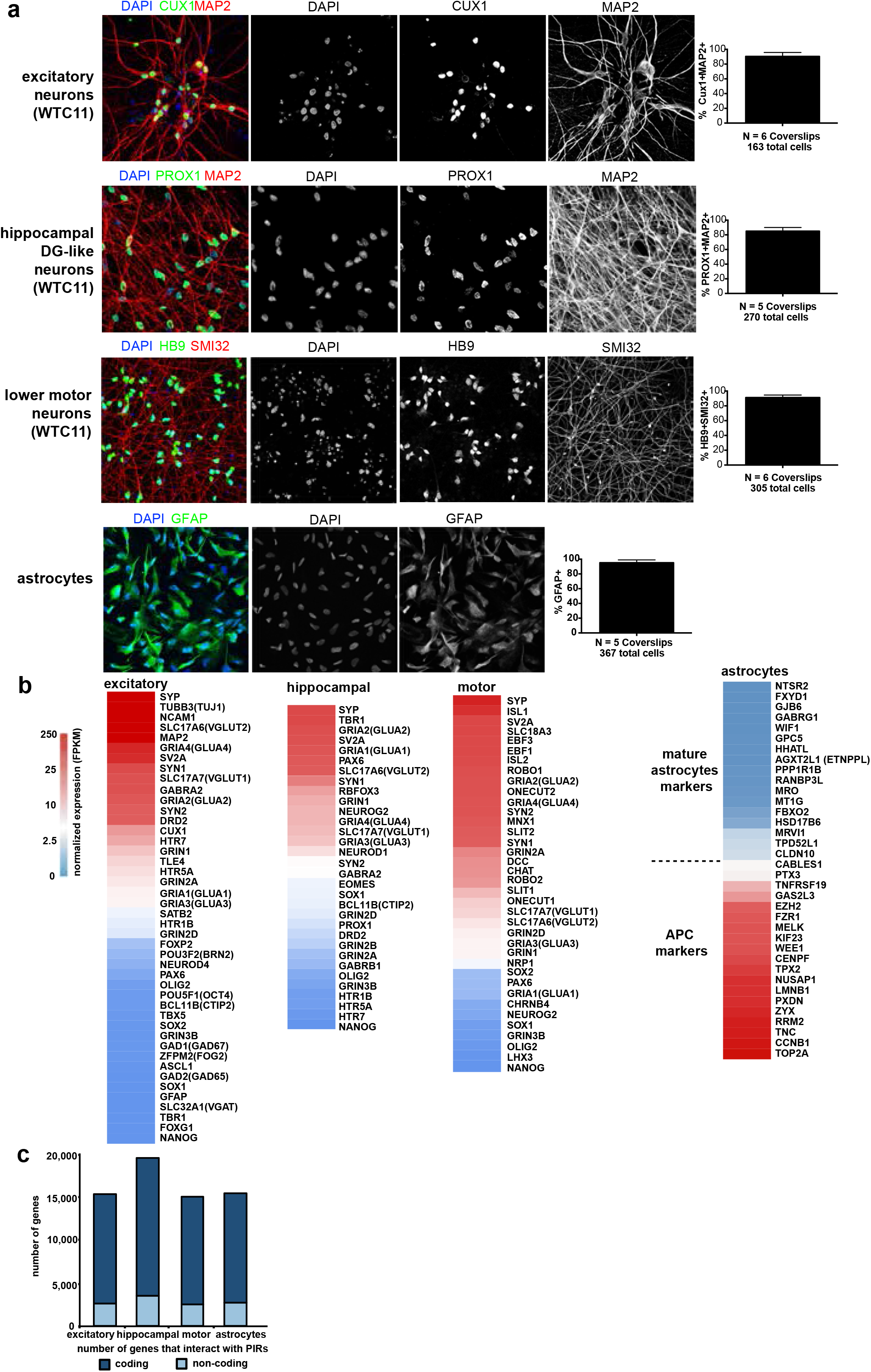
Characterization of the cell types used in the study. (**a**) Immunofluorescence staining of key markers in excitatory neurons, hippocampal DG-like neurons, lower motor neurons, and astrocytes. Excitatory neurons were positively stained for CUX1, an upper cortical layer marker, and MAP2, a neuronal marker which is specifically expressed in dendrites. The yield of excitatory neurons is calculated as the number of CUX1 and MAP2 double positive cells divided by the total number of live cells. Hippocampal DG-like neurons were positively stained for PROX1, a transcription factor specifying granule cell identity in the DG. The yield of mature hippocampal DG-like neurons is calculated as the number of PROX1 and MAP2 double positive cells divided by the total number of live cells. Lower motor neurons were positively stained for HB9, a motor neuron marker, and the pan-neuronal neurofilament marker SMI32. The yield of mature lower motor neurons is calculated as the number of HB9 and SMI32 double positive cells divided by the total number of live cells. Finally, astrocytes were positively stained for GFAP. The yield of GFAP-positive astrocytes is calculated as the number of GFAP positive cells divided by the total number of live cells. The number of staining experiments and the total number of cells is indicated, and error bars represent the SEM. (**b**) Heatmaps displaying the expression of key marker genes for the neural cell types. Astrocytes used in this study exhibit an expression profile consistent with APC identity. (**c**) Counts of protein coding (dark blue) and noncoding RNA (light blue) genes with promoters interacting in each cell type.

**Supplementary Figure 2.**
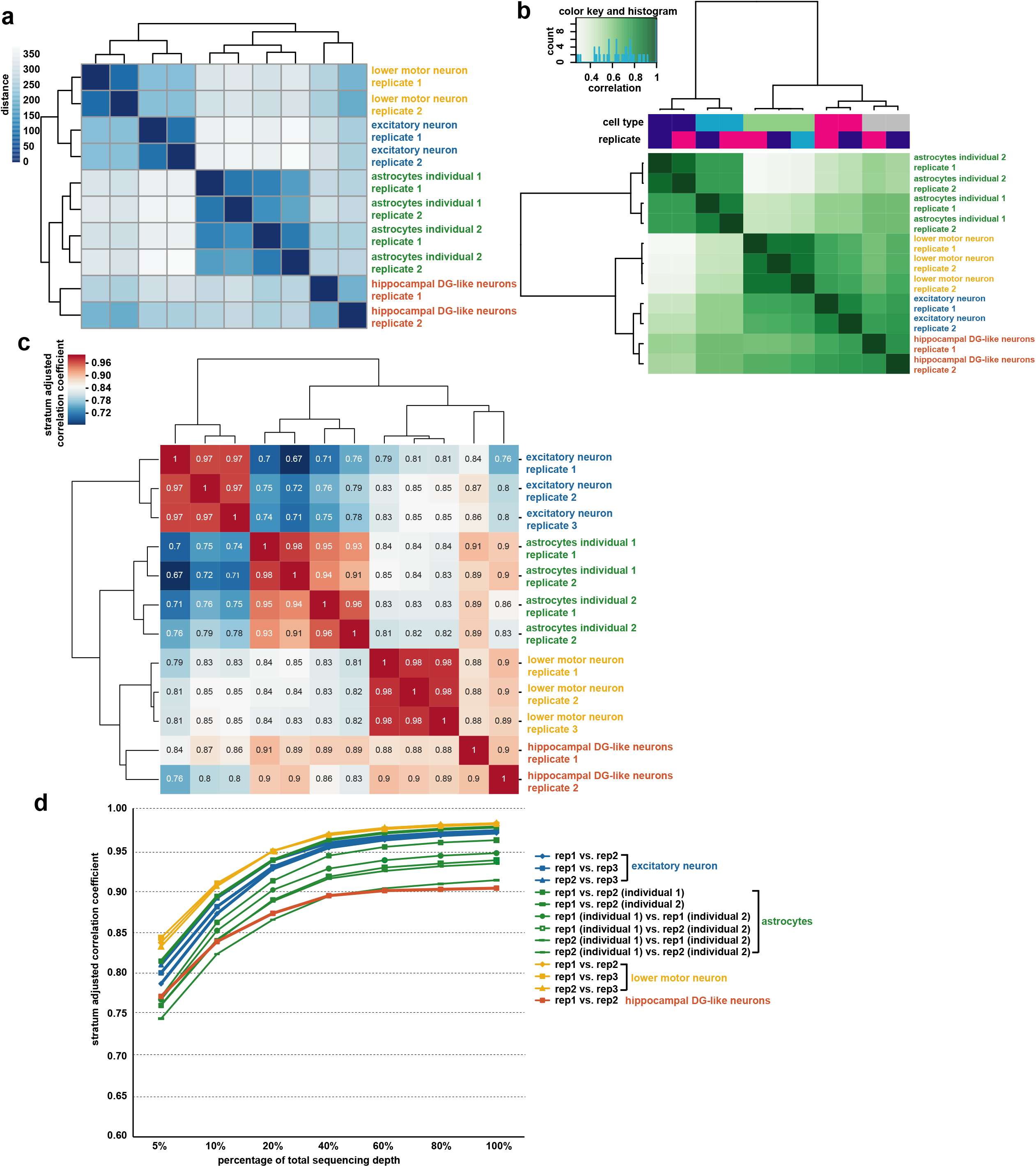
Correlation between pcHi-C, ATAC-seq, and RNA-seq replicates. (**a**) Gene expression values for each RNA-seq replicate were hierarchically clustered according to sample distances using DESeq2. (**b**) Heatmap with pairwise correlations and hierarchical clustering of read densities at the set of unified open chromatin peaks for the ATAC-seq replicates. (**c**) Heatmap with pairwise correlations based on the stratum-adjusted correlation coefficient (SCC) from HiCRep (evaluated at a resolution of 10 kb) for the pcHi-C replicates. (**d**) Saturation of the SCC between biological replicates for the pcHi-C libraries as a function of total sequencing depth.

**Supplementary Figure 3.**
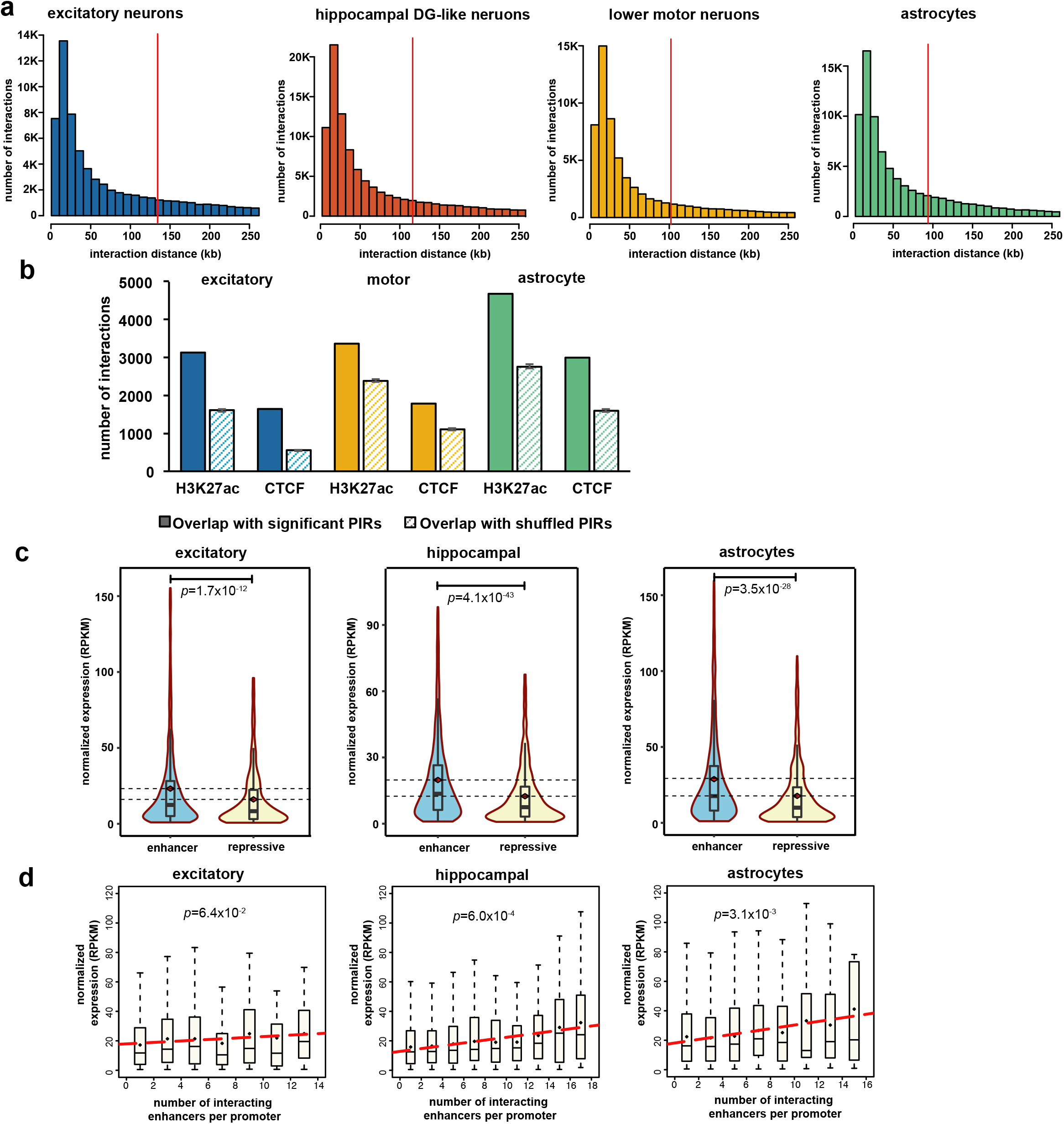
Integrative analysis of chromatin interactions in individual cell types. (**a**). Histograms of interaction distances for each cell type. The mean interaction distances for each cell type are indicated with red lines. (**b**) Bar plots showing counts of H3K27ac and CTCF binding sites overlapping significant (solid bars) versus randomly shuffled (striped bars) PIRs for excitatory neurons, lower motor neurons, and astrocytes. Error bars represent the standard deviation over 100 sampled sets of randomly shuffled PIRs. (**c**) Comparative gene expression analysis in individual cell types for expressed genes (normalized RPKM > 0.5) whose promoters interact exclusively with either enhancer-PIRs (n=6836) or repressive-PIRs (n=2612). Distributions of gene expression values are shown for each group. (**d**) Boxplots showing distributions of gene expression values in individual cell types for expressed genes (normalized RPKM > 0.5) grouped according to the numbers of interactions their promoters form with enhancer-PIRs. Linear regression was performed on the mean gene expression values for each bin. Only bins containing at least 10 genes were included in the analysis.

**Supplementary Figure 4.**
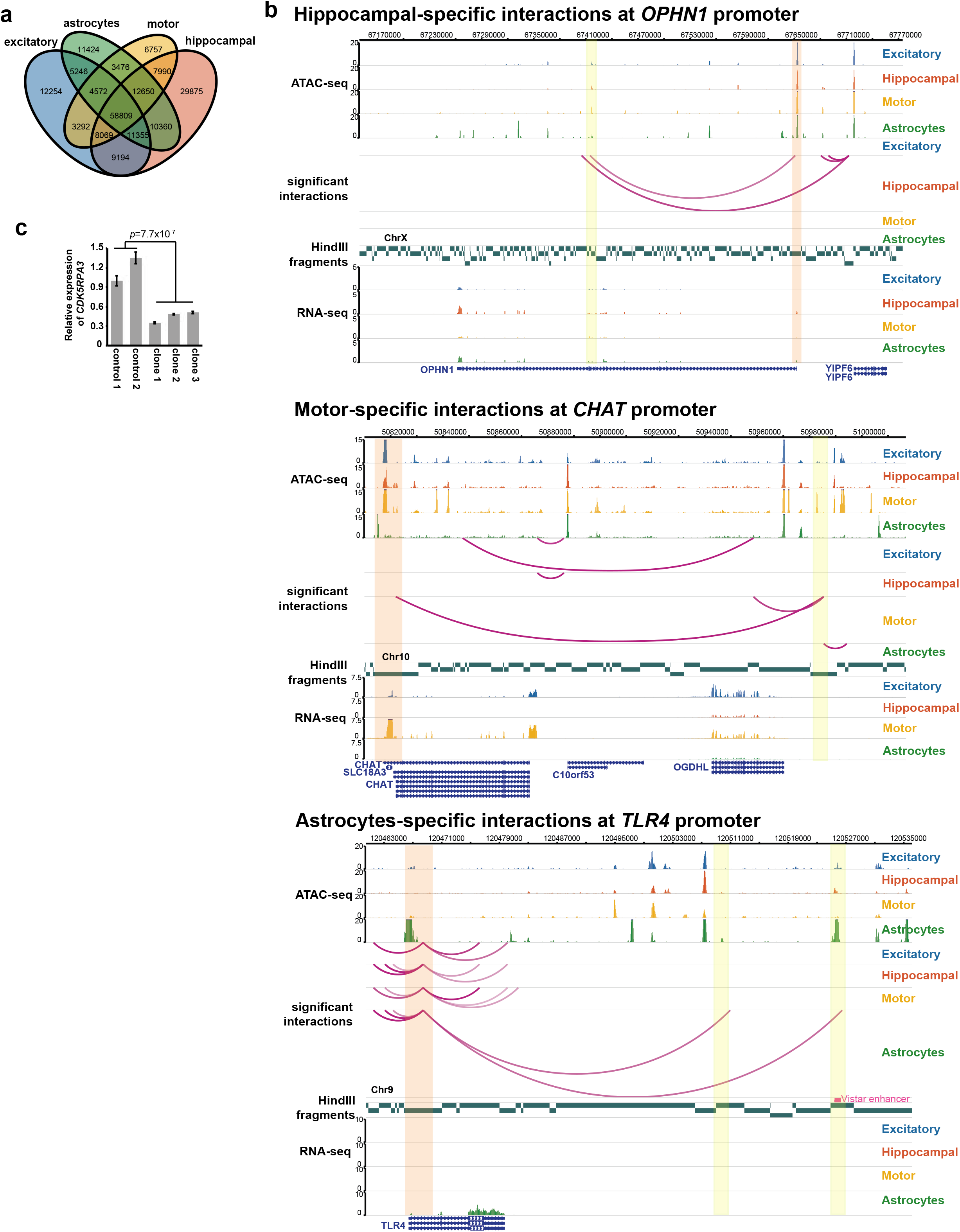
Cell type-specific aspects of chromatin interactions. (**a**) Venn diagram displaying counts of unique promoter-PIR interactions across excitatory neurons, hippocampal DG-like neurons, lower motor neurons, and astrocytes for each specificity pattern (groups 1-15 in **Fig. 3a**). (**b**) Examples of interactions between cell type-specific PIRs (yellow highlight) and the promoters for *OPHN1, CHAT*, and *TLR4* (orange highlight). Open chromatin peaks and gene expression are also displayed for each of the cell types. (**c**) Significant downregulation of *CDK5RAP5* expression was observed across three independent clones with homozygous deletions for the candidate PIR in excitatory neurons (two sample t-test, two-sided, *p*=7.7×10^−7^). Error bars represent the SEM.

**Supplementary Figure 5.**
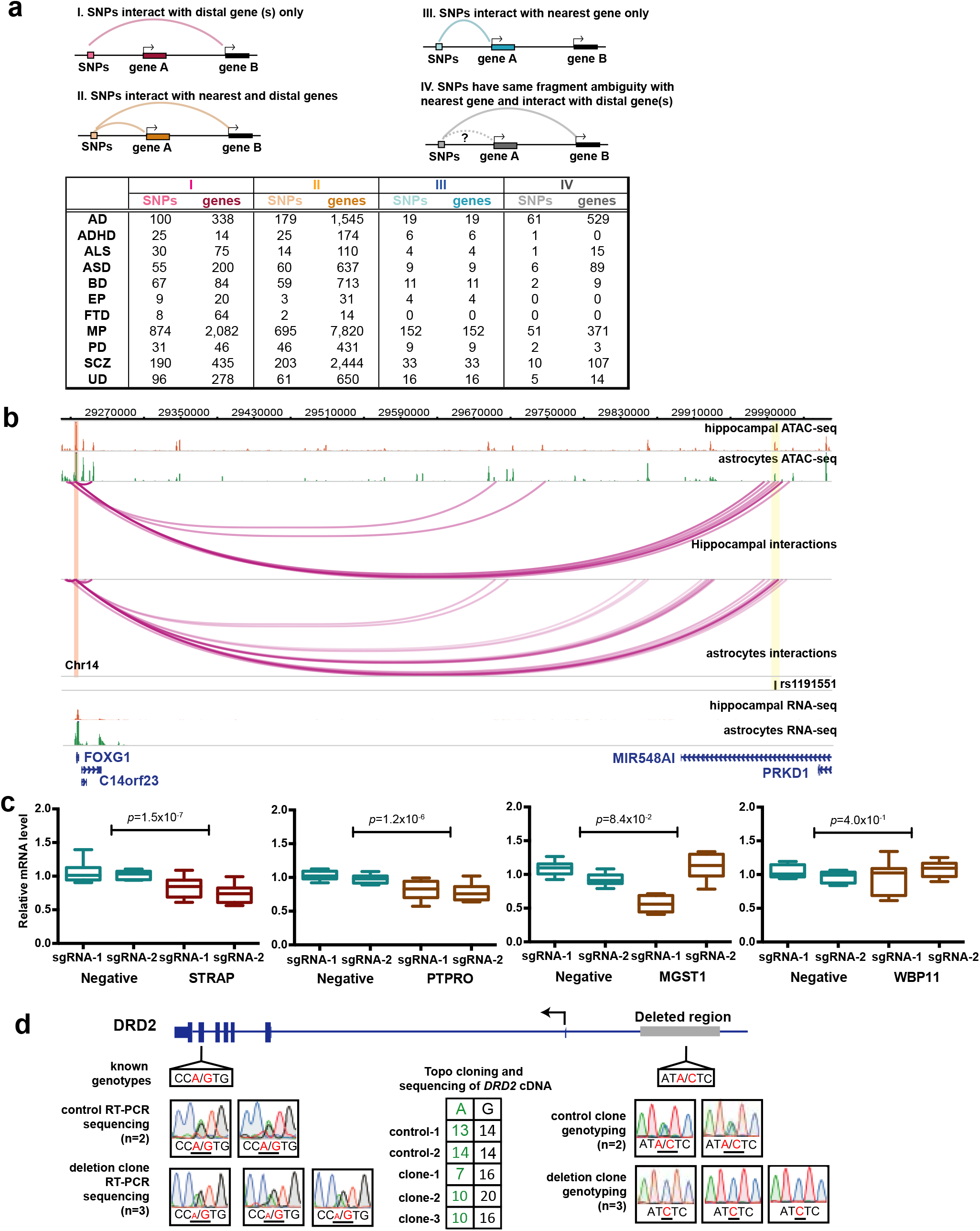
Using chromatin interactions to elucidate the functions of GWAS variants. (**a**) Counts of GWAS SNPs for each disease with at least one SNP in linkage interacting exclusively with their nearest genes (scenario III), interacting exclusively with more distal genes (scenario I), or interacting with both their nearest genes and more distal genes (scenario II). GWAS SNPs that could not be resolved for interactions with their nearest genes are also tabulated (scenario IV), along with counts of regulatory targets interacting with GWAS SNPs in each scenario. (**b**) Significant promoter-PIR interactions in hippocampal DG-like neurons and astrocytes recapitulate a previously reported interaction between the *FOXG1* promoter and a distal open chromatin peak containing rs1191551, a schizophrenia-associated variant^6^. (**c**) CRISPRi silencing of a candidate PIR for *STRAP* using two independent sgRNAs results in significant downregulation of *STRAP* expression in excitatory neurons (two sample t-test, two-sided, *p*=1.5×10^−7^). No significant downregulation was detected for the neighboring genes *MGST1* and *WBP1*, though the expression of *PTPRO* was affected. Each CRISPRi experiment was performed in triplicate, with three technical replicates per experiment. (**d**) Schematic of detected genotypes in the *DRD2* gene and its candidate PIR in mono-allelic deletion clones. Genotyping and RT-PCR sequencing for WTC11 variants in the *DRD2* gene reveal allele-specific imbalances in *DRD2* expression, consistent with the deletion of the candidate PIR in one of the alleles. Results for two wild type control clones and three mono-allelic deletion clones are shown for comparison.

**Supplementary Figure 6.**
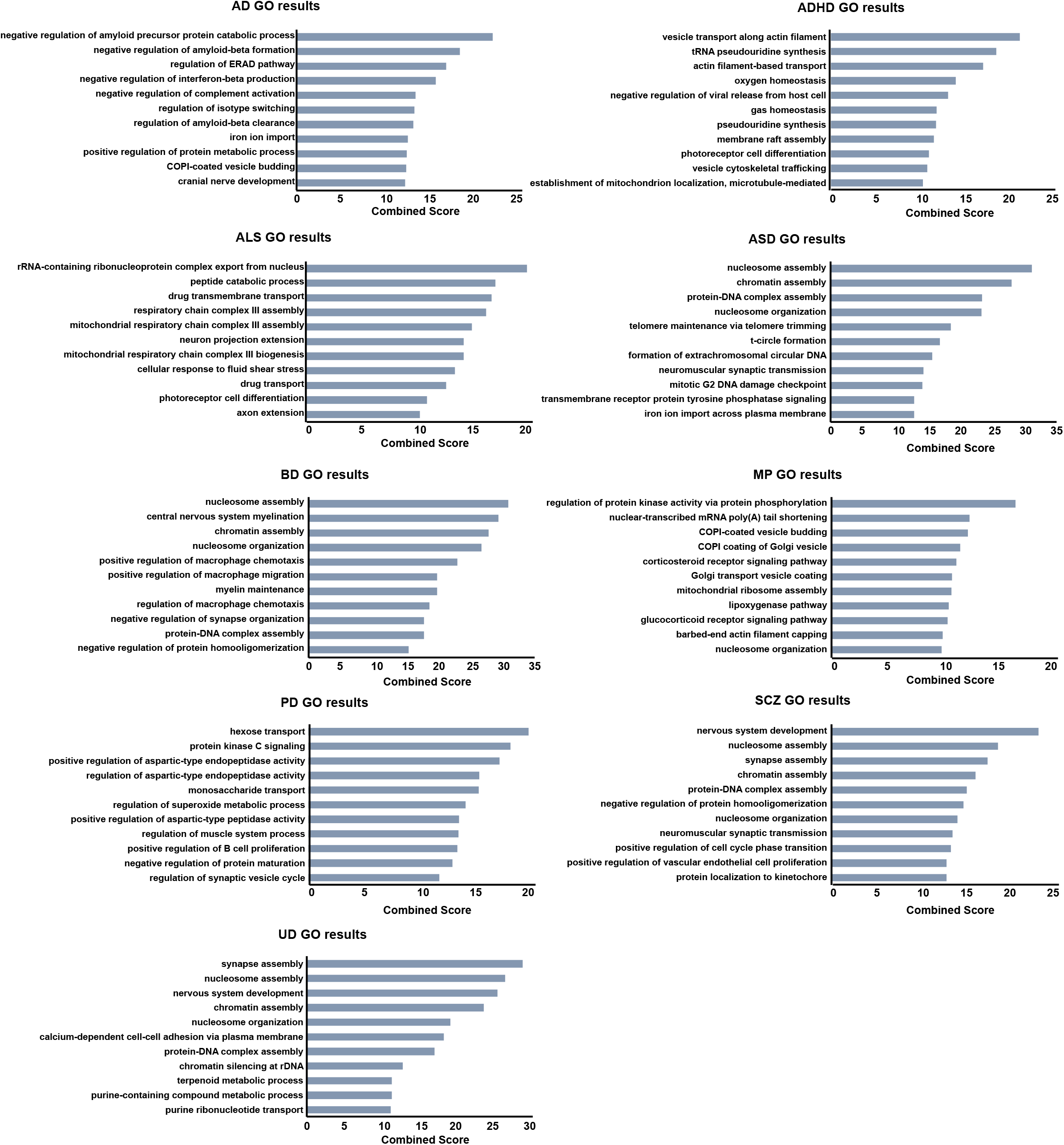
Top enriched GO terms for genes targeted by complex neurological disorder-associated variants. Top enriched GO terms from Enrichr for genes whose promoters are targeted by variants for each disease. EP and FTD are omitted due to their low numbers of reported variants and target genes identified by our significant promoter-PIR interactions. An expanded list of enriched GO terms is available in **Supplementary Table 9**. In general, we observe enrichment of terms associated with epigenetic, neuronal, and disease-specific processes across the diseases.

**Supplementary Figure 7.**
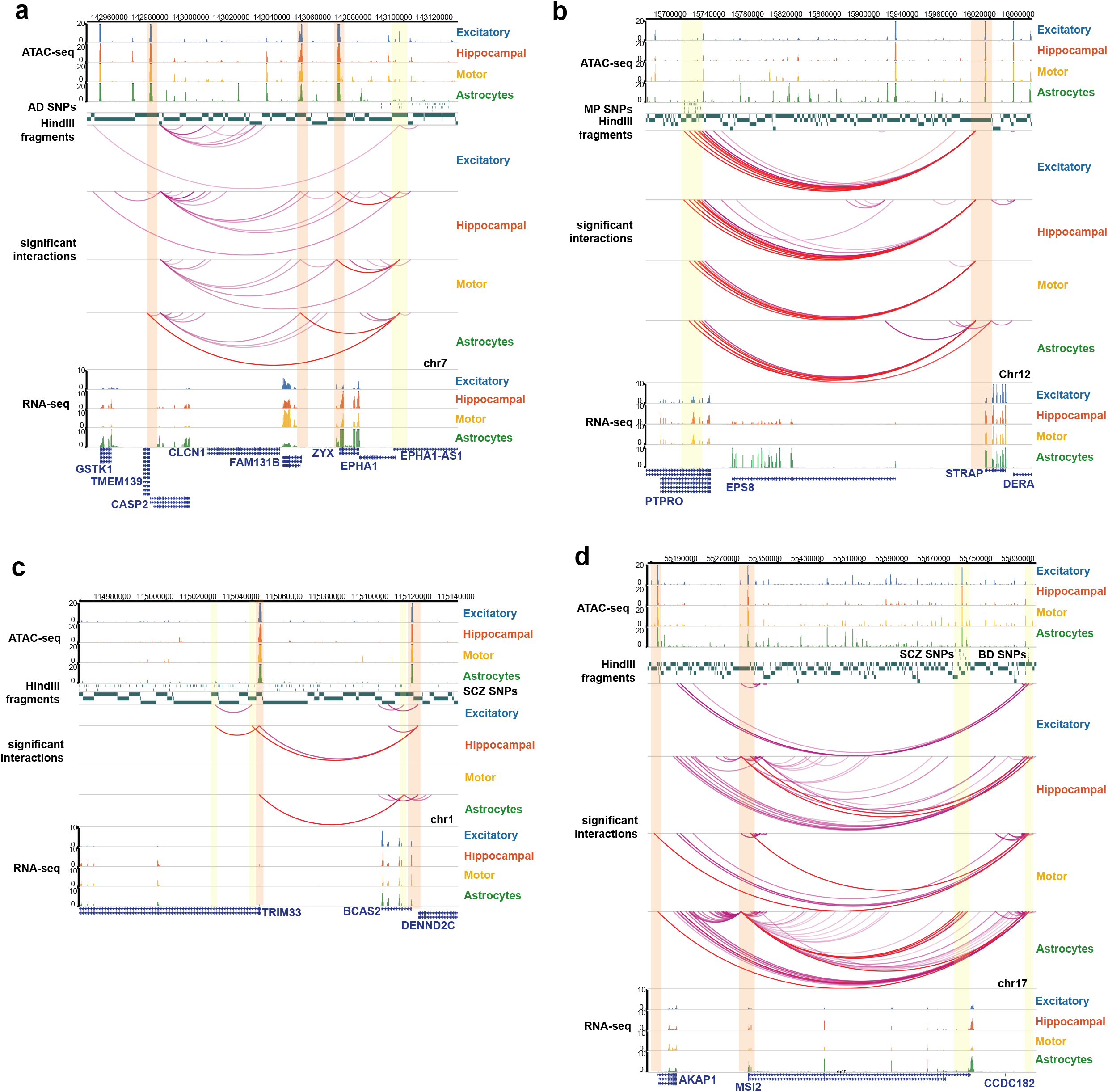
Examples of putative regulatory SNPs at cell type-specific PIRs. In all examples, interacting PIRs are highlighted in yellow and the targeted promoters are highlighted in orange. (**a**) A PIR containing AD SNPs interacts with the promoters of *FAM131B* and *CASP2* in astrocytes, but interacts instead with the *ZYX* promoter in hippocampal DG-like neurons and lower motor neurons. (**b**) PIRs containing MP SNPs in an intron for *PTPRO* interacts with the *STRAP* promoter in all four cell types. (**c**) A PIR containing SCZ SNPs interacts with the *TRIM33* promoter in astrocytes. Two other PIRs containing SCZ SNPs interact with the promoters of *TRIM33* and *BCAS2* in hippocampal DG-like neurons. (**d**) A PIR containing BD SNPs interacts with the *MSI2* promoter in hippocampal DG-like neurons, lower motor neurons, and astrocytes while also interacting with the *AKAP1* promoter in lower motor neurons and astrocytes. Meanwhile, another group of PIRs containing SCZ SNPs interact with the *MSI2* promoter exclusively in astrocytes.

## Tables

**Supplementary Table 1.** pcHi-C, ATAC-seq, and RNA-seq data processing metrics.

pcHi-C data processing metrics include output from the HiCUP mapping pipeline (columns C through O), the total number of reads in the CHiCAGO input file (columns P through Q), and the total number of processed interactions with score ≥ 5 from the CHiCAGO pipeline (column R). ATAC-seq data processing and QC metrics from the ENCODE pipeline are reported (columns C through K). The number of peaks called in individual as well as across all replicates is shown (column L). RNA-seq data processing metrics from STAR are reported (columns C through G).

**Supplementary Table 2.** Processed significant interactions called by CHiCAGO.

For each cell type, the left and right interacting BED intervals are shown in columns A through C and columns E through G, respectively. The number of supporting reads, interaction score, and specificity string for each interaction are shown in columns I through K. The number of overlaps with promoters, promoter open chromatin peaks, and distal open chromatin peaks, as well as the interacting gene IDs are shown in columns L through W. Overlaps for the left and right interacting BED intervals are shown separately, and “promoter” refers to protein coding and noncoding RNA transcripts in GENCODE 19 while “promoter other” refers to all other transcripts in GENCODE 19. The number of overlaps with positive Vista elements and SNPs for each disease and their associated IDs are shown separately for the left and right interacting BED intervals in columns X through BS.

**Supplementary Table 3.** GO enrichment results for genes interacting with cell type-specific PIRs.

GO enrichment results from the “Biological Process 2018” ontology in Enrichr are shown for genes interacting with PIRs specific to each of the cell types, as well as for genes interacting with PIRs shared across all the cell types (“shared terms”). In each tab, the top 100 GO terms and their associated p-values, Z-scores, combined scores, and genes are shown.

**Supplementary Table 4.** Motif enrichment results at cell type-specific PIRs.

For each cell type, the complete set of known motif results detected by HOMER are reported. This includes the motif name, consensus sequence, p-value, # of target sequences with motif, and # of background sequences with motif.

**Supplementary Table 5.** Putative target genes for *in vivo* validated enhancer elements.

Information about each positive Vista element including its position, ID, and annotation are shown in columns A through F. Information about its nearest gene, as well as genes whose promoters fall on the same HindIII fragment as the element are shown in columns G through I (protein coding and noncoding RNA transcripts in GENCODE 19 only). Column J represents all other transcripts from GENCODE 19 whose promoters fall on the same HindIII fragment as the element. Column K reports whether or not interactions are resolvable between the Vista element and its nearest gene in column G. Columns L through W contain information on whether the Vista element overlaps open chromatin peaks, participates in interactions, or targets genes on the other ends of interactions for each cell type.

**Supplementary Table 6.** GWAS Catalog SNP imputation summary.

The first tab contains a summary of the SNP imputation process for each disease in the study. This includes the number of GWAS SNPs downloaded from the GWAS Catalog, the number of GWAS SNPs passing the significance cutoff of 10^−6^, the numbers of GWAS SNPs associated with each study population, and the numbers of imputed SNPs for each study population. The remaining tabs contain lists of all the studies included for each disease along with their associated information.

**Supplementary Table 7.** Putative target genes for neurological disorder-associated SNPs.

Two tabs are included for each disease in Supplementary Table 7 (“GWAS SNPs” and “all SNPs”). The first tab (“GWAS SNPS”) contains the results for all GWAS SNPs downloaded from the GWAS Catalog. Information about each GWAS SNP including its position, rsid, allele information, query SNP status, and whether or not it overlaps any exons are shown in columns A through I. Information about its nearest gene, as well as genes whose promoters fall on the same HindIII fragment as the element are shown in columns J through L (protein coding and noncoding RNA transcripts in GENCODE 19 only). Column M represents all other transcripts from GENCODE 19 whose promoters fall on the same HindIII fragment as the element. Column N reports whether or not interactions are resolvable between the GWAS SNP (or any of its linked SNPs) and its nearest gene in column G. The total number of linked SNPs for each GWAS SNP is shown in column O. Columns P through AE contain information on whether the GWAS SNP itself or any of its linked SNPs participate in interactions or target genes on the other ends of interactions for each cell type. The second tab (“all SNPs”) contains similar information for all imputed SNPs. Columns A through I contain information about each imputed SNP, columns J through N contain information about its nearest and same fragment gene(s), and columns O through AD contain information on whether the imputed SNP overlaps with promoter or distal open chromatin peaks, participates in interactions, or targets genes on the other ends of interactions for each cell type.

**Supplementary Table 8.** Putative target genes for imputed SNPs overlapping open chromatin peaks. Supplementary Table 8 contains subsets of imputed SNPs from the “all SNPs” tabs in Supplementary Table 7, for which the imputed SNPs overlap with promoter or distal open chromatin peaks in at least one cell type.

**Supplementary Table 9.** GO enrichment results for disease-specific target genes.

GO enrichment results from the “Biological Process 2018” ontology in Enrichr are shown for genes interacting with PIRs containing variants for each disease. In each tab, the top 100 GO terms and their associated p-values, Z-scores, combined scores, and genes are shown.

**Supplementary Table 10.** Interactions exhibiting significant allelic bias.

A list of allelically biased interactions with a p-value cutoff of 10^−3^ are shown for the excitatory neurons and lower motor neurons. Supplementary Table 10 follows the format of Supplementary Table 2 with a few exceptions. The number of reads supporting interactions in each allele is reported in column I. The negative log p-value is reported in column J.

**Supplementary Table 11.** ASD epistatic SNP pairs by cell type.

The total numbers of epistatic SNP pairs in ASD cases and matched pseudo-controls with the following P-value thresholds: *p* ≤ 1.0×10^−20^, 1.0×10^−20^ ≤ *p* < 1.0×10^−17^, 1.0×10^−17^ ≤ *p* < 1.0×10^−14^, 1.0×10^−14^ ≤ *p* < 1.0×10^−13^, 1.0×10^−13^ ≤ *p* < 1.0×10^−12^, 1.0×10^−12^ ≤ *p* < 1.0×10^−11^, 1.0×10^−11^ ≤ *p* < 1.0×10^−10^, 1.0×10^−10^ ≤ *p* < 1.0×10^−9^, 1.0×10^−9^ ≤ *p* < 1.0×10^−8^, and 1.0×10^−8^ ≤ *p* ≤ 1.0×10^−7^ are shown for the entire dataset including for excitatory neurons, hippocampal DG-like neurons, lower motor neurons, and astrocytes. Results for nonsignificant interactions in the corresponding cell types are also shown.

**Supplementary Table 12.** sgRNA and primer sequences.

A full list of sgRNAs and primers for the CRISPR deletion and CRISPRi experiments are reported here.

